# Structural mechanism of MUC5AC mucin net-like polymer formation and its SNP variability that affect risk of the lung diseases COPD and IPF

**DOI:** 10.1101/2024.08.02.606332

**Authors:** Sergio Trillo-Muyo, Anna Ermund, Jesper M. Magnusson, Gunnar C. Hansson

## Abstract

Gel-forming mucins MUC5AC and MUC5B constitute the main structural component of the mucus in the respiratory system. Secreted mucins interact specifically with each other and other molecules giving mucus specific properties. We determined the cryoEM structures of the wild type MUC5AC-D3 assembly and the structural SNP variants R996Q and R1201W. Our structures explain the basis of MUC5AC N-terminal non-covalent oligomerization upon secretion. The MUC5AC-D3 assembly forms covalent dimers in two alternative conformations, open and closed. The closed conformation dimers interact through an arginine rich loop in the TIL3 domain forming tetramers. Moreover, we found a positive disease correlation between the SNP (R996Q, rs878913005), Chronic Obstructive Pulmonary Disease (COPD), and Idiopathic Pulmonary Fibrosis (IPF). The well-known MUC5B promotor SNP (rs35705950) association with IPF is much stronger when combined with the MUC5AC SNP. Our study provides a model to explain the formation of MUC5AC net-like structures and how both SNPs will affect mucus organization and increase risk of lung disease.

## INTRODUCTION

Mucins are a group of highly glycosylated molecules that cover and protect all mucosal surfaces of the body (Hansson, 2020). The two gel-forming mucins MUC5AC and MUC5B constitute the main structural components of the mucus protecting the underlying epithelia in the respiratory system. Both mucins share the same domain organization and are large proteins, 5,654 or 5,762 amino acids and a mass of 586 or 596 kDa without the characteristic *O*-glycosylation. They are related to the other gel-forming mucins MUC2 and MUC6 and the von Willebrand factor (VWF). All these mucins have an N-terminal part built by 3.5 von Willebrand D assemblies, VWD1, VWD2, VWD’ and VWD3. The VWD assemblies are formed by the VWD domain, C8, trypsin inhibitor like (TIL) and E domains, except the VWD’ domain that only has the TIL and E domains. The gel-forming mucins have their N-termini followed by one or several PTS domains rich in proline, threonine and serine, often in a repetitive fashion. The hydroxy amino acids in the PTS domains become heavily *O*-glycosylated to form the extended rod-like mucin domains. The dense sugar coating is responsible for the mucin hygroscopicity necessary for binding water and to mucus formation. The PTS domains are typically interrupted by a variable number of CysD domains, nine in MUC5AC and seven in MUC5B, involved in homotypic interactions as shown for CysD2 of MUC2 (Recktenwald *et al*., 2024). The MUC5AC, MUC5B and MUC2 mucin C-terminal region is formed by a VWD4 assembly, 3.5 VWC domains and a C-terminal cysteine-knot (Gallego *et al*., 2023).

Secreted mucins interact specifically with each other and other molecules giving mucus specific properties. These mucins are shown to be orderly packed in the goblet cell granulke due to low pH and high Ca^2+^. Upon secretion, they are unpacked into large disulfide-based polymers. The process requires an increase in pH and maybe calcium removal, both of which can be reached by bicarbonate transported via the cystic fibrosis transmembrane conductance regulator (CFTR). The inter-molecular disulfide bonds formed in the ER and trans-Golgi network in the C-terminus and the N-terminal regions pave the ground for the expansion into covalent linear oligomers. However, additional non-covalent interactions are required for mucus organization.

The respiratory system is constantly exposed to inhaled particles, bacteria and viruses. Humans and pigs have submucosal glands down to 10^th^ branch. They secrete a chloride and bicarbonate rich fluid that pulls out the MUC5B mucin into long polymers exiting the glands as >20 µm thick bundled strands (Ermund *et al*., 2017;Hoegger *et al*., 2014;Widdicombe and Wine, 2015). These contains >1,000 parallel MUC5B molecules that interacting laterally. These are patchily coated by the MUC5AC from the surface goblet cells which controls the bundle movement by attachment/detachment events that together with MUC5AC threads clean the larger airways (Bos *et al*., 2023;Ermund *et al*., 2018;Ermund *et al*., 2017).

The MUC5B mucin is required for the normal lung homeostasis in mice whereas the MUC5AC is not (Roy *et al*., 2014). Interestingly, the MUC5AC mucin is induced and increased in amount at metaplasia and diseases of the lung (Fahy and Dickey, 2010;Fernandez-Blanco *et al*., 2018). This affects the surface goblet cells at metaplasia to co-express both MUC5B and MUC5AC in the same cell and even in the same granule (Hoang *et al*., 2022), leading to the formation of an attached stratified mucus layer in the airways (Fernandez-Blanco *et al*., 2018).

The question why we have two different mucins in the lung and in what way they differ as been a long-time puzzle. The MUCB mucin is clearly forming linear molecules which was confirmed by electron microscopy (Ermund *et al*., 2017;Ridley *et al*., 2016). The MUC5AC mucin on the other hand forms net-like structures (Carpenter *et al*., 2021). We have now addressed the question why MUC5AC is forming a differently structured mucus by structural studies using cryoEM. The results show that the disulfide bonded linear MUC5AC mucin also interacts non-covalently in its VWD3 assembly and that there are genetic variants in this region that are linked to increased risk of lung diseases.

## RESULTS

### MUC5AC-N covalent dimer

Expression of the secreted complete N-terminal part of MUC5AC (MUC5AC-N) has proved to be challenging and has only been achieved at low yields in polarized airway cell lines (Ryan *et al*., 2015). We have tried to produce the complete secreted MUC5AC in CHO and HEK cells, also without any success. The VWD1-VWD2 (D1-D2) assemblies of VWF and MUC2 are cleaved off after polymerization suggesting that these domains are required for packing, but less important for the mature protein (Sadler, 1998;Schutte *et al*., 2014). MUC2 D1-D2 assemblies are compactly packed at low pH in the goblet cell granule at the same time as they are involved in intracellular polymerization (Javitt *et al*., 2020). MUC2-N expressed in CHO cells and analyzed by cryoEM at pH 7.4 shows that only the compact D3 dimer assembly can be visualized, the D1, D2 and D’ are too flexible (unpublished). This suggests less importance of D1-D2 for the mature secreted mucins at physiological, neutral pH. We therefore focused on the MUC5AC-D3 (Figure 1A and B) assembly and its interactions.

**Figure 1.**
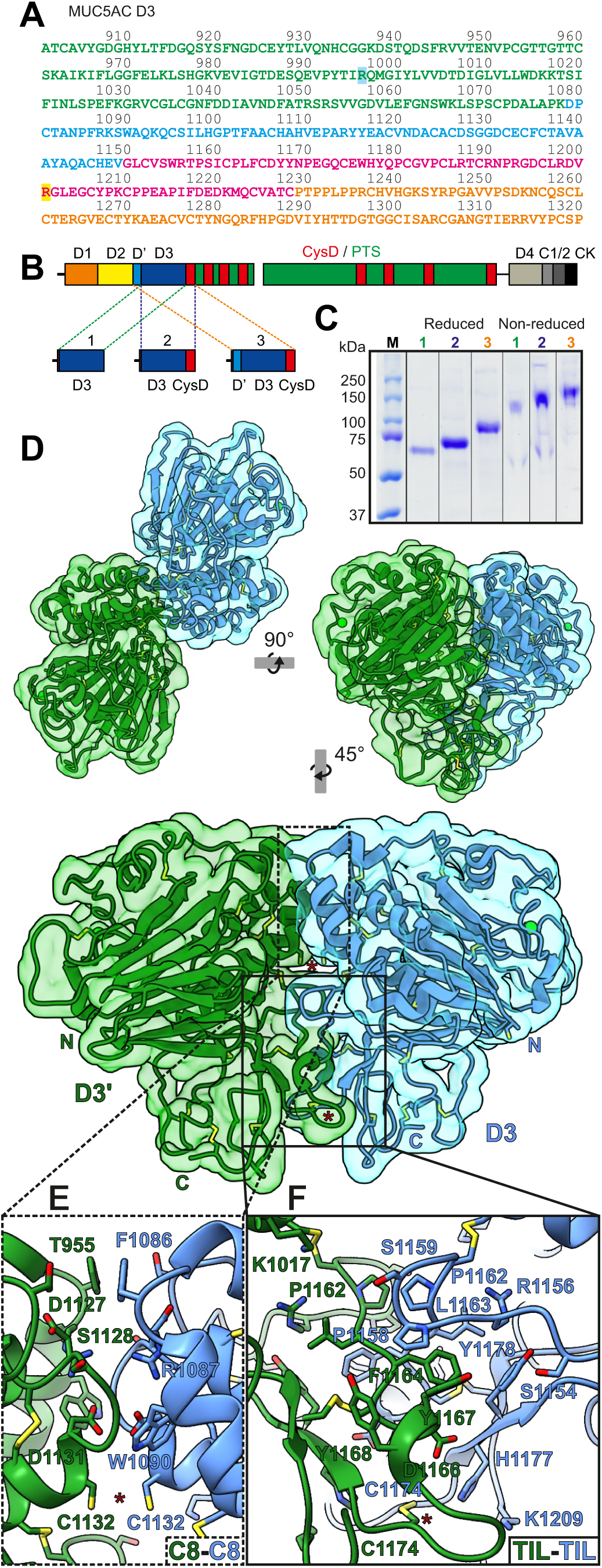
CryoEM structure of the MUC5AC-D3 assembly. (A) MUC5AC-D3 sequence. VWD3 sequence is showed in green, C8-3 in blue, TIL3 in pink and E3 in orange. The residues affected by the SNPs rs36189285 (R996) and rs878913005 (R1201) are highlighted in blue and yellow respectively. (B) Schematic sketch of the domains of MUC5AC mucin with the N-terminal region (VWD1 (orange), VWD2 (yellow), VWD’ (light blue) and VWD3 (dark blue)), nine CysD domains (red) surrounded by PTS sequences densely *O*-glycosylated to form mucin domains (green) and the C-terminal region (VWD4 (light grey), VWCs (dark grey) and CK (black)). The fragments analyzed are enlarged. (C) SDS-PAGE analysis of reduced and non-reduced D3 (1), D3-CysD (2) and D’-D3-CysD (3) reveals the formation of reducible dimers in all three fragments. (D) MUC5AC D3 assembly cryoEM density map and cartoon representation showing the disulfide bonds. The map and model of the two monomers are shown in green and cyan. The Ca^2+^ ions are shown as green spheres. The top left figure represents the top view of the molecule. It is rotated anticlockwise by 90°around the x-axis in the top right figure, and rotated clockwise by 45°around the y-axis and enlarged by 50% in the bottom figure showing the details of the front view. Putative intermolecular disulfide bonds are marked by red starts (Cys1132-Cys1132’ bond seems to be reduced). N-terminal (N) and C-terminal (C) of each monomer are marked. (E) Detail of the MUC5AC-D3 covalent dimerization interface zoomed in from (D) showing the interaction between C8-3 domains. (F) Detail of the MUC5AC-D3 covalent dimerization interface zoomed in from (D) showing the TIL3-TIL3’ interaction.

We designed expression plasmids including the N-terminal D’ assembly, the D3 assembly and the CysD1 domain with and without an N-terminal 6xHis-tag (Figure 1B). These plasmids were expressed in Lec 3.2.8.1 CHO cells and the recombinant proteins purified and analyzed by gel electrophoresis showing the expected band sizes after reduction (Figure 1C). Without reduction, the bands migrate approximately at the double size, suggesting that all three recombinant proteins are disulfide bonded dimers.

The structure of the MUC5AC-D3 assembly was analyzed by cryoEM at pH 7.4 (Table S1, Figure S1 and S2). The cryoEM reconstruction of the D’-D3-CysD1 recombinant protein reached higher resolution (3.2 Å) than the isolated D3, even if only the D3 assembly is visible due to flexibility of the D’ and CysD1 domains (Figure 1D). The overall structure of the MUC5AC-D3 assembly dimer is similar to the previously reported D3 assemblies (Dong *et al*., 2019;Javitt *et al*., 2019). The covalent MUC5AC-D3 assembly dimer is formed via two intermolecular disulfide bonds, Cys1132-Cys1132 in the C8-3 domain and Cys1174-Cys1174 in the TIL3 domain (Figure 1D-F). However, the Cys1132 inter-molecular disulfide bond is partially reduced as observed previously in MUC2-D3 (Javitt *et al*., 2019). The interaction surface is highly conserved between MUC5AC-D3 and MUC2-D3 with only three relevant substitutions. In the C8-3 domain interaction region MUC5AC-D3 presents a phenylalanine (Phe1086, Figure 1E) instead of a histidine (MUC2-D3 His1042), potentially affecting the effect of pH in the intracellular packing. In the TIL3 domain interface, Ser1154 (Figure 1F) substitutes a phenylalanine present in an equivalent position in MUC2 (Phe1110). This serine is highly conserved in MUC5AC and MUC5B between species. The hydrophobic pocket formed by Phe1164’, Tyr1167’ and Tyr1168’ interacts with Pro1158, while Tyr1178, Leu1163 and the aliphatic chain of Arg1156 creates another pocket that accommodates Phe1164’ (Figure 1F). In MUC2 the second hydrophobic pocket is also formed by Phe1110 stabilizing the interaction with Phe1164’. In the same area, MUC2 interfacing residues Arg973’-Asp1115 are substituted in MUC5AC by Ser1159-Lys1017’ resulting in a loss of a salt bridge. No hydrogen bonds between Ser1159-Lys1017’ were observed.

The structure of the MUC2-E3 domain could be solved by crystallography due to the crystal packaging contacts showing high B-factors (Javitt *et al*., 2019). However, in solution MUC5AC is too flexible to be modeled. After 3D refinement only some noisy densities appeared in the area where it should be located.

### MUC5AC-D3 TIL3 structure

The overall structure of MUC5AC (Figure 2A) is similar to MUC2 and VWF. However, an overlay of MUC5AC and MUC2 shows differences including the loops β4-β5 and β8-β9 in VWD3 domain and the loop β1-β2 in the TIL3 domain, all located on the same side of the molecule (Figure 2B). The TIL3 domain is the region of the molecule presenting more differences when compared with MUC2-D3 and VWF-D3 assemblies (Figure 2C). The MUC5AC-TIL3 β1-β2 loop, unlike the other mucins and VWF, is remarkably rich in arginines (20%) (Figure 2D). The disulfide bond organization in this domain is unique but still shows characteristics in common with the structures previously described. The disulfide bonds between MUC5AC-TIL3 cysteines 1165-1206, 1189-1228 and 1210-1224 are conserved at similar positions in all three proteins, in addition to the Cys1174 forming the intermolecular dimer disulfide bond. The Cys1181 interacts with the C8-3 region in MUC5AC and MUC2 (Cys1137) stabilizing the domain organization while the equivalent cysteine in VWF (Cys1149) is involved in an extra interdomain disulfide bond. On the other hand, MUC5AC and VWF have an identical disulfide bond in the center of the loop β1-β2 that is absent in MUC2, Cys1185-Cys1196 in MUC5AC and Cys1153-Cys1165 in VWF. MUC2 contains an *N*-glycosylation site in this area, Asn1154, not present in the other mucins or VWF. The MUC5B-TIL3 domain shares 74.2% identity with MUC5AC and almost certainly displays the same disulfide bond arrangement.

**Figure 2.**
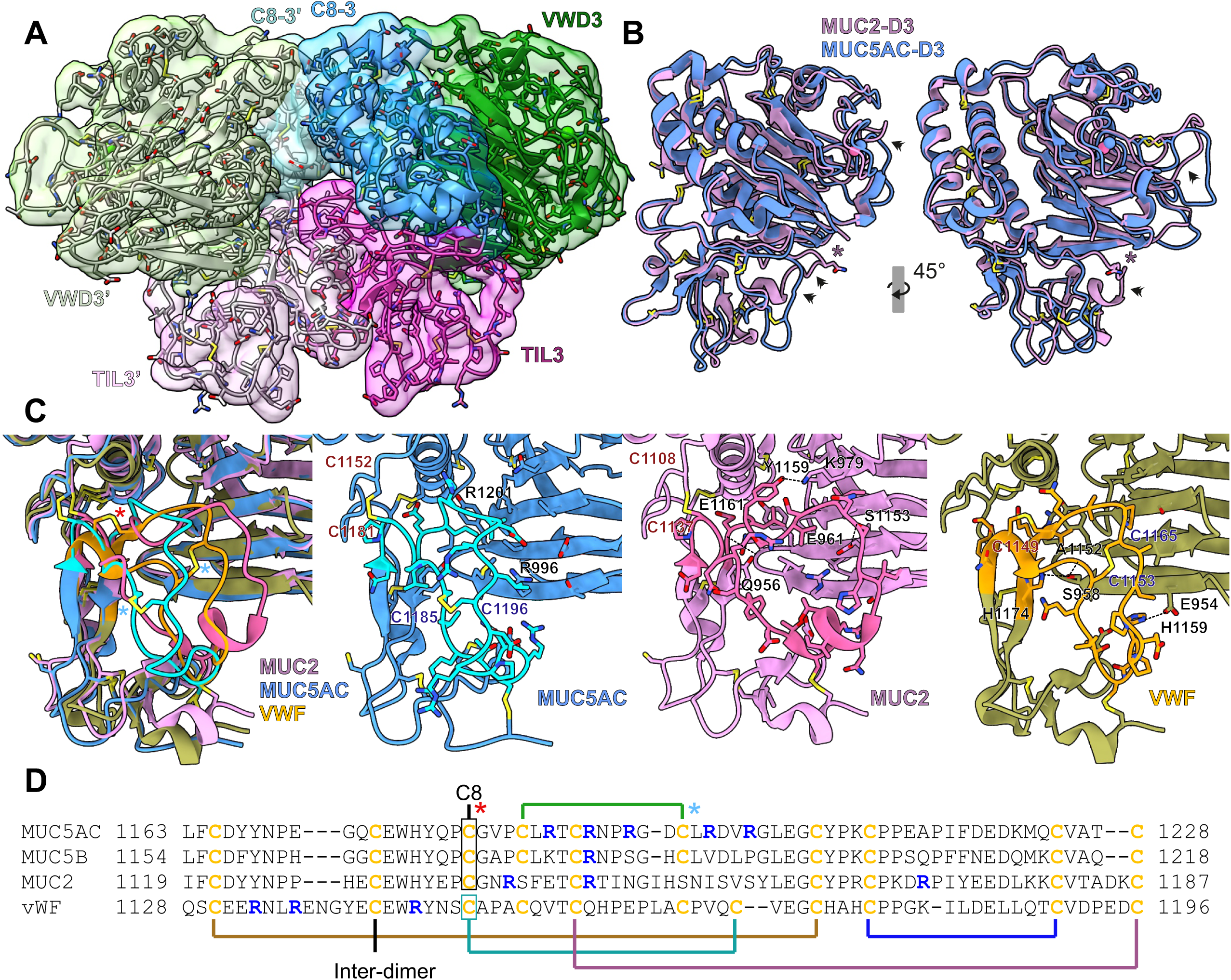
MUC5AC-D3 domains. (A) Front view of MUC5AC-D3 assembly dimer from cryoEM density map and model. The two VWD3 domains are shown in dark and light green, C8-3 domains in dark and light blue, and TIL3 domains in dark and light pink. (B) Structural alignment of MUC5AC-D3 (blue) and MUC2-D3 (pink. PDB code: 6rbf). Left image is presented in the same orientation as (A). Right image is rotated clockwise 45°. Regions with high variability are marked by black arrows. The *N*-glycosylated Asn1154 in lateral chain of MUC2 is marked with a black star. (C) Detail of TIL3 structurally aligned of MUC5AC-D3 (blue), MUC2-D3 (pink. PDB code: 6rbf) and VWF-D3 (orange. PDB code: 6n29). The TIL3 β1-β2 loop is highlighted in brighter colors. To the left, superposition of all three structures showing the larger distance between the TIL3 and VWD3 domains in MUC5AC. The distinct disulfide bonds are marked by stars, in red the one connecting the TIL3 domain with C8-3 domain and in blue the internal TIL3 β1-β2 loop disulfide bond. All three structures are shown separately showing the TIL3 β1-β2 loop and interfacing residues of lateral chains. The cysteines involved in the distinctive disulfide pattern are labeled. Hydrogen bonds between VWD3 and TIL3 are showed with dashed black lines and the residues involved are annotated. In MUC5AC, the residues affected by SNP variation at the amino acids R996 and R1201 are marked. (D) Amino acid sequence alignment of MUC5AC, MUC5B, MUC2 and VWF TIL3 domains. Disulfide bonds are marked. Stars mark distinct disulfide bonds as in (C). Cysteines are colored yellow and arginines in blue.

There are also remarkable differences in the relative position of the whole TIL3 domain in relation to VWD3. In MUC2 and VWF, the center of the TIL3 β1-β2 loop is located only 6.2 Å and 8.8 Å away from VWD3 β9, while in MUC5AC they are separated by 14.4 Å. This close contact is explained by the presence of multiple hydrogen bonds between these domains in MUC2 (Gln956-Glu1161, Glu961-Ser1153 and Lys979-Tyr1159) and VWF (Glu954-His1159, Ser958-Ala1152 and Ser958-His1174) that are completely absent in MUC5AC.

### MUC5AC-D3 open conformation

During the 3D-classification processes, a second MUC5AC-D3 assembly conformation was observed. The 3D volume refined with these particles showed low resolution with a severe preferred orientation problem. The particles were 2D classified showing all classes having the same top view orientation (Figure 3A). The maximum diameter of the new open conformation classes is 30% higher than in the standard or closed conformation. The 2D top view of both conformations display enough details to recognize the VWD3 domain β-sandwich and the C8-3 domain, and together with the low-resolution 3D map allowed us to model this conformation into the density maps (Figure 3B). The VWD-C8 interaction dissociates and exposes a hydrophobic surface in VWD3 to the solvent (Figure 3C). The domains establish a novel interaction between the loops β7-β8 and β11-β12 in VWD3 and the C-terminal side of α4 in C8-3, and between the VWD3 β8 and β1-β2 TIL3 loop (Figure 3D). The long connecting loop between VWD3 and C8-3 domains makes this conformational change possible (Figure 3B). This conformational change could explain the previous observation that MUC5AC binds significantly more to hydrophobic surfaces compared to MUC5B (Carpenter *et al*., 2021).

**Figure 3.**
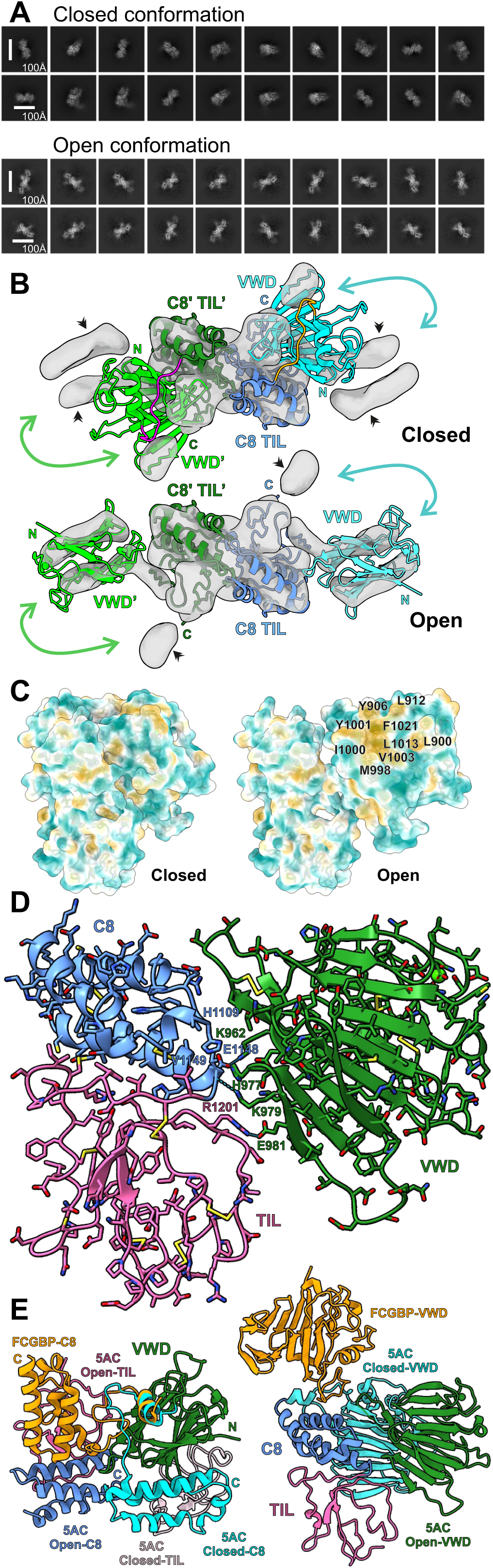
MUC5AC-D3 assembly open conformation. (A) CryoEM 2D classes, box size 220Å. The top figure shows the closed conformation 2D classes from the high-resolution structure shown in Figure 1 and 2. The discarded particles from an initial 3D classification were further 2D classified. These 2D classes are shown in the bottom panel, open conformation. (B) CryoEM low-resolution map generated using the particles from (A) bottom panel. The top figure shows the fitting on the closed conformation model and the bottom the proposed model for the open conformation. The VWD3 domain of one monomer is shown in light green, C8-3 and TIL3 domains in dark blue and the connecting loop in magenta. The VWD3 domain of the other monomer is shown in cyan, C8-3 and TIL3 in blue and the connecting loop in orange. The black arrows show the densities not covered by the models. The non-occupied densities in the closed form are explained by the movement of VWD3 as shown by the cyan arrows. The C-terminal of TIL3 points toward the marked densities in the open conformation as they could represent E3 and/or CysD. (C) Surface representation of the closed (left) and open (right) conformation colored by molecular lipophilicity potential (MPL) from dark cyan (most hydrophilic) via white to dark goldenrod (most lipophilic). The newly exposed hydrophobic pocket residues in the open conformation are labeled. (D) Detail of proposed model for MUC5AC-D3 open conformation. VWD3 is colored in green, C8-3 in blue and TIL3 in pink. Putative salt bridges between K962 and E1148, and E981 and R1201, and the hydrogen bond between K979 and V1149 are represented by dashed lines. These residues and the interfacing histidines His977 and His1109 are labeled. (E) MUC5AC-D3 closed and open conformation and FCGBP D10 alignment (Yeshaya *et al*., 2024). In the left figure MUC5AC-D3 closed (C8-3 in cyan and TIL3 in light pink) and open (C8-3 in blue and TIL3 in dark pink) conformation and FCGBP D10 (C8-10 in orange) were aligned to the VWD domain (green). The N-terminal (N) of VWD and C-terminal (C) of the different C8 domains are marked following the same color code. In the right figure MUC5AC-D3 closed (cyan) and open (green) conformation and FCGBP D10 (orange) were aligned by the C8 (blue) and TIL (pink) domains.

Recently an open structure from the D10 assembly in FCGBP has been published (Yeshaya *et al*., 2024), supporting the idea that the opening of the VWD assemblies could have a physiological function. However, there are multiple differences between these two structures (Figure 3E). The FCGBP structure was solved by crystallography and therefore the crystal packaging could promote or alter the structure of the open conformation. The relative position of the domains is completely different. In the MUC5AC-D3 closed conformation the N-terminal of VWD and the C-terminal of C8 points in the same direction and are only 12 Å apart. In FCGBP they are 55 Å apart and rotated 90°, while in MUC5AC open conformation they are at 34Å and rotated -45°. Furthermore, in FCGBP the domains interact only through the connecting loop as there are no other intramolecular interactions in contrast to MUC5AC (Figure 3D). The FCGBP-D10 structure lacks the TIL domain known to interact with the VWD domain in other VWD assemblies to maintain the compact structure where the conformational changes were explained by the GDPH autocatalytic cleavage not present in MUC5AC-D3 (Yeshaya *et al*., 2024).

### MUC5AC-D3 variants

Interestingly, when genomic databases of MUC5AC were analyzed, the parts of MUC5AC most different from the other mucins as discussed above (Figure 2) contained two variants where a single amino acid had been replaced (Figure 4A). In both of these, one of the arginines typical for MUC5AC was replaced by a glutamine (Arg996Gln, rs36189285) or by tryptophan (Arg1201Trp, rs878913005). We produced recombinant proteins with each variant separate as well as with both arginines replaced. The structure of each of these were solved by cryoEM and compared (Table S1; Figures S3-S6). The general appearance of the dimers is essentially identical (Figure 4B). However, careful comparison of each variant to the WT elucidates the effects of the mutations (Figure 4A). The Arg996Gln mutation, located in VWD3 β9, is placed directly in the interface between the VWD3 and the TIL3 domain. The substitution results in a slight reduction of the distance between the two domains (Figure 4A). No interactions involving this residue are observed in the WT or in Arg996Gln, but considering the resolution limitations a hydrogen bond between Arg996 or Gln996 and a TIL3 β1-β2 loop residue cannot be completely ruled out. The observed alteration can also be attributed to arginine steric repulsion.

**Figure 4.**
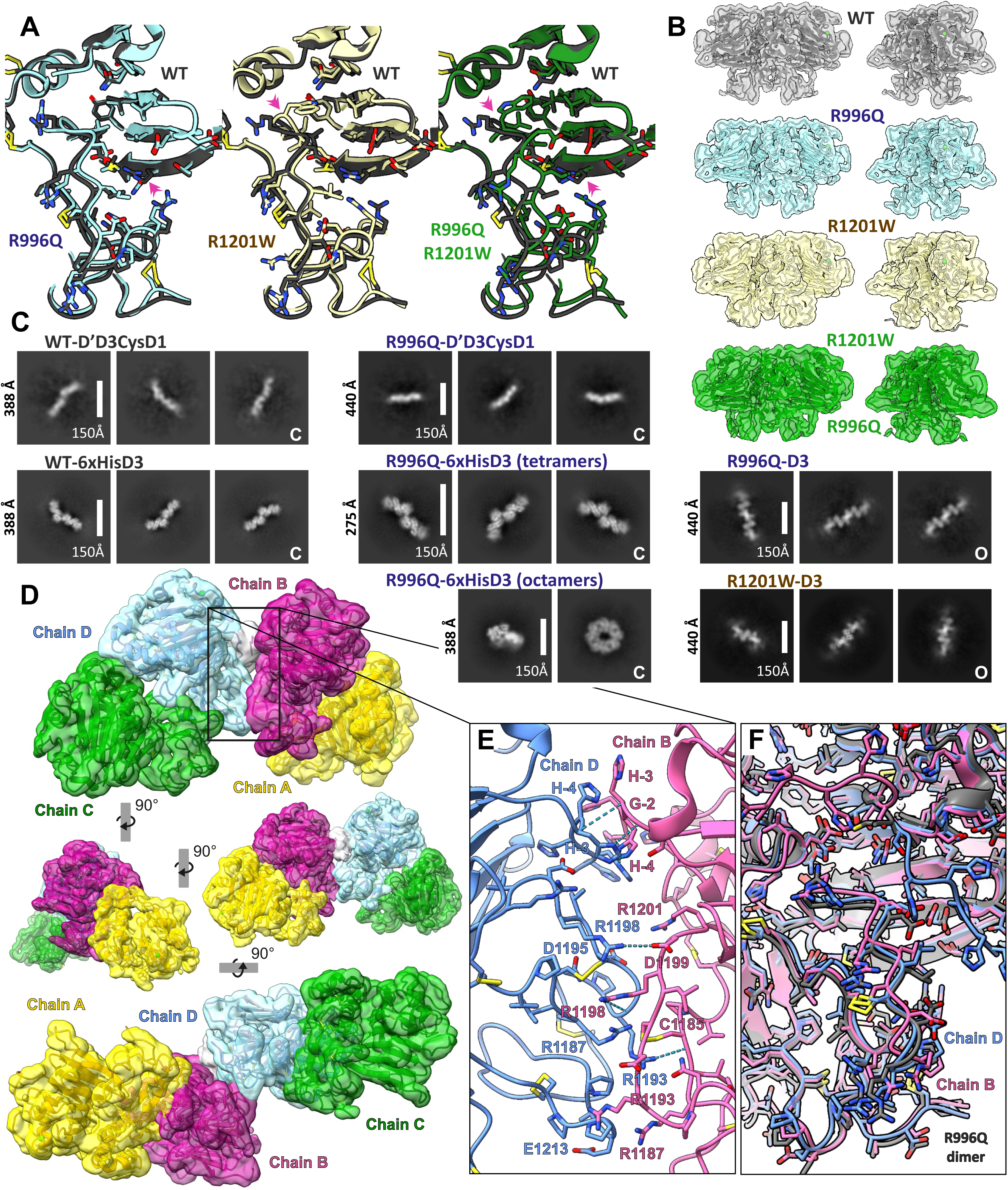
MUC5AC-D3 tetramerization. (A) Detail of TIL3-VWD3 interface mutants aligned with the MUC5AC-D3 WT (dark grey). The left figure shows the Arg996Gln mutant in cyan, the mid shows the Arg1201Trp mutant in yellow, and the right shows the double mutant Arg996Gln-Arg1201Trp in green. The mutations are pointed by pink arrows. (B) CryoEM density map and cartoon representation of the MUC5AC-D3 dimeric assemblies WT (grey), Arg996Gln (cyan), Arg1201Trp (yellow) and Arg996Gln-Arg1201Trp (green) at two 45° angles. (C) CryoEM 2D classes of higher order oligomers. Box sizes are specified for every group of classes. The groups of closed conformation oligomers are marked with “C” and the open conformation with “O”. (D) MUC5AC-D3 Arg996Gln tetrameric assembly cryoEM density map and cartoon representation. One covalent dimer is shown in yellow (chain A) and magenta (chain B) and the other in green (chain C) and cyan (chain D). The His-tag density is shown in white. The top image shows a lateral view of the tetramer. It is rotated clockwise by 90° around y-axis and reduced 1.5 times in the middle-left figure, rotated clockwise by 90° around y-axis again in the middle-right figure, and rotated anticlockwise by 90°around x-axis and rescaled to the original size in the bottom figure. (E) Detail of the MUC5AC-D3 non-covalent tetramerization interface zoomed from (D). The predicted salt bridges and hydrogens bonds are shown as dashed cyan lines. (F) Structural alignment of the tetramer chain B (pink) and chain D (blue) against the R996Q dimer chain A (grey).

In contrast, the Arg1201Trp mutation, placed in the β1-β2 TIL3 loop, only affects the adjacent residues. The solvent exposed arginine is substituted by a tryptophan pointing toward the VWD-C8/TIL interface. The tryptophan accommodates into the VWD3 hydrophobic interfacing region in close proximity to Leu1013. In the double mutant, the effect of the Arg996Gln mutation is more pronounced, bringing the VWD3 and the TIL3 domain closest together. Even though no major rearrangements were found, both SNPs can affect the equilibrium between the dimer open and closed conformation. Open conformation dimers were constantly found during cryoEM 2D and 3D classifications in the WT assembly. Only when 2D templates from WT open conformation were used for particle picking, these could be detected in the Arg996Gln sample.

### MUC5AC-D3 tetramers

The grids of the MUC5AC-D’D3-CysD1 assembly showed the covalent dimers (Figure 2), but also numerous images showed tetrameric particles (Figure 4C). In the same way, all MUC5AC variants showed some tetramers in cryoEM. The interaction is shown to be flexible as most of the 2D classes are noisy and 3D reconstructions only show elongated blobs, but the presence of the His-tag stabilizes it. The WT 6xHis-MUC5AC-D3 assembly shows 2D classes with structural features just in one orientation and the 3D reconstructions thus have low quality. Similarly, the structure of the tetrameric 6xHis-MUC5AC-D3 Arg996Gln assembly was solved at low resolution (6-7 Å), not enough to reveal the molecular mechanism of the non-covalent oligomerization. Surprisingly, the 6xHis-MUC5AC-D3 Arg996Gln adopts another conformation not observed in the other variants as the protein forms ring shape octamers (Figure 4C). The ring formation gives stability to the interactions. However, after refinement only one of the tetramers was well defined, the others were poorly defined due to remaining flexibilities and difficulties in isolating octamers from tetramers in some orientations. Particle subtraction followed by local refinement let us solve the structure of a tetramer forming part of the octamer at 3.69 Å resolution (Table S1; Figures 4 D and E and S6).

The MUC5AC-D3 non-covalent tetramer interaction is driven by the TIL3 domain and involves an interface area of 385.4 Å^2^ (Figure 4D). The interaction between the MUC5AC TIL3 domains occurs in the arginine-rich loop β1-β2 stabilized by a disulfide bond between Cys1185-Cys1196 (Figure 2C and D). The interaction is mainly hydrophilic (Figure 4E). Intermolecular salt bridges and hydrogen bonds are formed between lateral chains of Asp1199-chainB (Asp1199B) and Arg1198-chainD (Arg1198D), and between Arg1198B lateral chain and the main chain of Arg1193D and Asp1195D, Arg1193D and Cys1185B, and between Arg1198D and Leu1197B and Arg1198B. The model also shows the presence of the lateral chain of Arg1187D and Arg1193D in the proximity of the Cys1185B-Cys1196B disulfide bond and Arg1198B pointing to the same disulfide bond in chain D.

The comparison between the tetrameric MUC5AC-D3 Arg996Gln assembly and the free closed conformation dimer shows significant differences only in the TIL3 domain region (Figure 4F). While the tetramer chain D shows only minor deviations, the TIL3 loop β1-β2 in chain B changes its relative position with C8-3 and VWD3 domains allowing the interaction with chain D.

The solved structure confirms the role of the His-tag in the interaction. No interactions between the His-tag of one monomer and the core of the other one were observed, just His-tag– His-tag interactions, supporting that it is an unspecific interaction that stabilizes the TIL3-TIL3 interface (Figure 4E).

Moreover, 2D classifications suggest that the MUC5AC-D3 assembly can also form an additional kind of tetramers involving the open conformation (Figure 4C). This interaction is linear and leads to the formation of high-order oligomers that look longest in Arg996Gln. However, the open conformation oligomers were only found in assemblies lacking D’ or N-terminal 6xHis-tag. The 2D classes together with the open conformation low-resolution map (Figure 3B) suggested a major steric impediment at the N-terminal region. The symmetric interaction of two VWD3 domains through the external side of the β-sheet 1 seems to bury both N-terminal residues. Assemblies containing D’ or N-terminal His-tag were only found to form tetramers based in closed conformation covalent dimers, making it likely that the open form tetramers are an artefact.

### MUC5AC-D3 tetramerization and physiological properties

The tetrameric MUC5AC will cross-link linear molecules with an angle of about 40° as illustrated in Figure 5A. The *O*-glycosylated PTS domains are extended rods and thus expected to drive the MUC5AC N-termini apart. This will generate a net-like structure as suggested in its ideal form in Figure 5B. This model is well in line with what we know of MUC5AC organization from staining of tissues. When tissue sections of the surface mucus of the stomach was stained for MUC5AC, a stratified and laminated organization appears (Figure 5C). This is similar to the inner mucus layer of the colon made up by the MUC2 mucin. Electron microscopy of the tracheal surface shows the different organization of the two lung mucins, linear bundled strands of MUC5B and more net-like appearances of the MUC5AC (Figure 5D). The different organization of these two mucins is as shown by Carpenter *et al*. when pure MUC5B and MUC5AC mucins were studies by EM (Carpenter *et al*., 2021).

**Figure 5.**
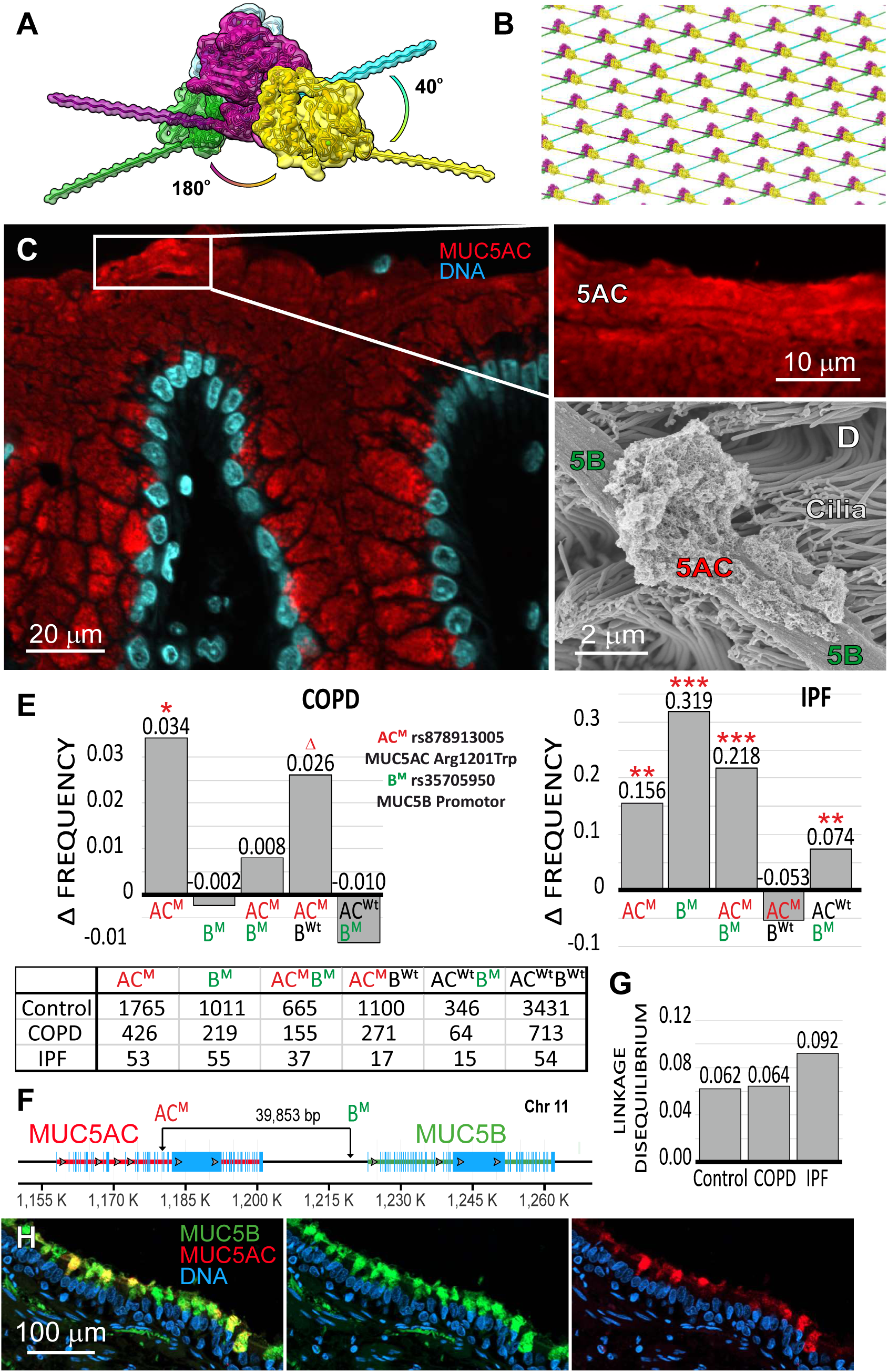
MUC5AC-D3 physiological properties, SNPs and correlation with disease. (A) MUC5AC-D3 Arg996Gln tetrameric assembly cryoEM density map and cartoon representation. One covalent dimer is shown in yellow (chain A) and magenta (chain B) and the other in green (chain C) and cyan (chain D). The PTS domains are schematically represented as a polyalanine straighten chain protruding from the TIL3 C-termini. The PTS from each covalent dimer extend in opposite directions forming an angle of about 40° with the PTS from the other dimer. (B) Ideal schematic representation of the MUC5AC network generated by repetitions of (A) linked by covalent dimerization at the end of the PTS domains (cystine-knot domain). (C) Carnoy fixed human stomach biopsy paraffin section stained with a monoclonal anti-human MUC5AC antibody (45M1; red) and Hoechst (nuclei; blue). The enlarged white square shows stratified surface mucus positive for MUC5AC. (D) Scanning electron micrograph of a piglet airway showing a MUC5B bundled strand, MUC5AC mucus attached to the bundle, and cilia. (E) Frequency of SNPs increases in COPD (left) and IPF (right). AC^M^ stands for mutant MUC5AC Arg1201Trp (rs878913005), B^M^ for mutant MUC5B promotor (rs35705950), AC^Wt^ for wild type in MUC5AC Arg1201 position and B^Wt^ for wild type MUC5B promotor in the position for rs35705950. Significance with the Fisher exact test is shown by three stars (p<0.001), one star (p<0.05) or a triangle (p<0.1). The bottom table shows the raw values used for the frequency calculations. (F) Linkage disequilibrium between MUC5AC Arg1201Trp (rs878913005) and MUC5B promotor (rs35705950) in (E) control, COPD and IPF groups. The graphic shows the frequency increase of both mutations appearing in the same subject in relation to the expected frequency if both mutations were independent. (G) Genomic organization of the *MUC5AC* and *MUC5B* gene locus och chromosome 11. (H) Formalin fixed paraffin section from an IPF lung explanted at lung transplantation, stained with a polyclonal anti-human MUC5B antibody (green), a monoclonal anti-human MUC5AC antibody (45M1; red) and Hoechst (nuclei; blue).

### MUC5AC-D3 SNPs and disease

The SNP sequence variant where Arg1201 is replaced by Trp (rs878913005) has not normally been included in GWAS studies as it is located close to the variable region of the MUC5AC mucin. To analyze if there is any coupling of this SNP to disease, the complete genome of individuals of the UK-biobank were analyzed for this SNP and compared to available medical information. The SNP (Arg1201Trp) was found to be overrepresented in patients with COPD (n=1,139). The frequency is 0.3740 in the COPD cohort while it is 0.3397 in the control group (n=5,196), representing a statistically significant increase at p<0.05 (Fisher exact test statistic value 0.0278) of 0.0343 (10%) (AC^M^ in Figure 5E). This SNP was also analyzed for patients with idiopathic pulmonary fibrosis (IPF). Although this cohort was relatively small (n=107), an increased frequency rise of 0.1556 (46%) was observed at statistical significance (p<0.01, Fisher exact test statistic value 0.0013).

In order to discard the influence of other known SNPs in IPF, the well documented rs35705950 SNP affecting the MUC5B mucin promotor region was analyzed in the UK-biobank individuals. As expected, a significant (p<0.001) increase in frequency of this SNP in IPF was found for rs35705950, 0.3194 (164%) (B^M^ in Figure 5E). Interestingly, when both SNPs were analyzed in relation to each other, we observed that the association between SNPs and disease is much stronger when both SNPs appear at the same time. The increased frequency of presenting the two SNPs at the same time in IPF patients is three times higher than when only the rs35705950 in MUC5B promotor combined with WT for MUC5AC, 0.2180 p<0.001 and 0.0736 p<0.01 respectively (Figure 5E). In contrast, the effect of the MUC5AC SNP rs878913005 on the frequency of COPD is not related to the rs35705950 MUC5B.

The MUC5AC and MUC5B mucins are located after each other on chromosome 11 with 39,853 nucleotides in between (Figure 5F). A genetic link between the two is thus possible. Analysis for linkage disequilibrium revealed a value of 0.062 and 0.064 between rs878913005 (MUC5AC) and rs35705950 (MUC5B) in the control and COPD groups, respectively. However, the linkage disequilibrium between the two SNPs increased up to 0.092 (48%) in the IPF cohort (Figure 5G). Consequently, the frequency of having both SNPs increases from the expected 0.066 to 0.128 (0.066+0.062) in the control group while it increases from 0.254 to 0.346 (0.254+0.092) in IPF, meaning that the probability of presenting both SNPs is almost 3 times higher in IPF patients. Together the results suggest that the MUC5AC variant (Arg1201Trp) show increased risk of COPD, but the risk for IPF is particularly increased.

The analysis of the other MUC5AC SNP (rs36189285, Arg996Gln) studied here, was inconclusive due to its low frequency in the European population and therefore in the UK-biobank. This SNP is more frequent in the Asian population.

IPF is characterized by fibrosis in the smaller airways where the MUC5B mucin is normally predominant (Okuda *et al*., 2018). When peripheral human lung tissue from patients with IPF were stained, most surface secretory cells expressed both the MUC5B and MUC5AC mucins (Figure 5H) further suggesting that the MUC5AC could have a role in IPF.

## DISCUSSION

The D3 mediated covalent dimerization in the MUC5AC mucin described in this study as predicted by homology with MUC2, together with the C-terminal dimerization observed in all gel-forming mucins, lead to the formation of long linear polymers (Gallego *et al*., 2023;Javitt *et al*., 2020). Different assemblies and structural rearrangements occur along the secretory pathway to achieve an efficient unpacking upon release and thus the N and the C-terminal dimerization must occur in an ordered way. First, the C-terminal inter disulfide bond is formed in the ER (pH 7.4) and then in the trans-Golgi network (pH 6.2) the N-termini are coupled together (Asker *et al*., 1998;Axelsson *et al*., 1998). To prevent the N-terminal disulfide bond formation in the ER, the interaction of the covalent oligomerization interface should only happen at lower pH. The dominance of charged residues close to the interface could have a role regulating the formation of the covalent link along the secretory pathway, especially His1177 and Asp1166, conserved in all gel-forming mucins. Different structures of the MUC2-D3 assembly has been solved by crystallography and cryoEM, all of them at low pH (Javitt *et al*., 2019;Javitt *et al*., 2020). We present the first mucin D3 assembly solved at neutral pH, but interestingly no major structural differences were observed. It indicates that once the covalent dimerization is stabilized the interface is locked, even if the Cys1132-Cys1132 dimer bond is partially reduced. This disulfide bond reduction was first observed for the equivalent Cys observed in the MUC2 crystallographic structure. This was attributed to radiation damage, an event that could also occur in cryoEM and can provide information about “weak links" that can be of structural significance (Weik *et al*., 2000). This suggests that the formation of the Cys1132-Cys1132 disulfide bond could be tightly regulated by its redox potential, requiring the higher oxidizing environment found in the Golgi compared to the ER (Kellokumpu, 2019). Its higher radiation damage susceptibility could also be explained by the formation of a stabilizing S⋯O interaction (Bhattacharyya *et al*., 2020) with the carbonyl *O*-atom from the same cysteine located just at 3.3 Å from the *S*-atom.

Nevertheless, observation of MUC5AC under physiological conditions has shown that it is different from MUC5B in that it does not appear as long linear polymers (Carpenter *et al*., 2021;Ermund *et al*., 2017;Ridley *et al*., 2016). Instead, it forms more complex and likely net-like structures as shown here and previously (Carpenter *et al*., 2021;Ryan *et al*., 2015), requiring the existence of other intermolecular interactions. The here observed capability of MUC5AC to form non-covalent dimers through the TIL3 domain and thus MUC5AC-D3 tetramers explains the molecular mechanism for the net-like polymer formation. The interaction is mainly hydrophilic as it is based on multiple hydrogen bonds and salt bridges through a densely positively charged region. Unlike the covalent MUC5AC-D3 dimerization that locks the assembly, the non-covalent formation of tetramers is flexible and can be regulated by pH and ionic strength upon secretion. The unique MUC5AC arginine rich TIL3 composition suggests that this interaction only occurs in MUC5AC.

MUC5AC can appear both in a closed and an open conformation. Although the open model is derived from low resolution cryoEM, it shows that the VWD assemblies can open in solution supporting the physiological relevance of the previously reported FCGBP VWD10 structure (Yeshaya *et al*., 2024). Considering other known VWD3 assemblies, we cannot exclude the existence of an open VWF or MUC2-D3 open conformation, but the closer interactions between the TIL domain and VWD make them less energetically favorable. The open conformation implies the exposure of a highly hydrophobic surface that is not evident in FCGBP-D10. The likely artefactual open conformation tetramers and high order oligomers found in MUC5AC-D3 suggest that the external site of the VWD3 β-sheet 1 becomes a new interaction surface that could establish contacts with other mucins, mucus associated proteins, or other domains from the same molecule. This could explain the hydrophobic properties attributed to the mucins but do not explained any of the actual structures.

In addition to the observed non-covalent interactions within the VWD3 assembly explaining the net-like appearance of MUC5AC, there are additional possible interactions within mucins. The CysD2 domain of MUC2 was recently shown to form weak homotypic dimers that was further stabilized by transglutamination catalyzed by TGM3 (Recktenwald *et al*., 2024). Human MUC5AC contains nine CysD domains, thus similar mechanisms could take place also in MUC5AC to produce a highly crosslinked mucin. However, although all the CysD domains have a common structure, there is a large variation in especially surface amino acids opening for variation in the interactions.

SNPs affecting the respiratory gel-forming mucins have been previously associated with disease (Sabo *et al*., 2023;Seibold *et al*., 2011;Shrine *et al*., 2019). However, mucin sequencing has proved to be challenging due to their highly repetitive sequences in the PTS domain (Svensson *et al*., 2018). Mucin sequences and SNPs have traditionally been poorly annotated, especially in these domains and flanking regions, including the VWD3 assembly. Here we observed that two frequent missense SNPs in the MUC5AC-D3 assembly, rs36189285 and rs878913005, are located in the interaction surfaces involved in the MUC5AC tetramerization. The rs36189285 SNP corresponds to Arg996Gln, while rs878913005 corresponds to Arg1201Trp, both substituting the for MUC5AC typical arginines. Arg996Gln is present in 32.9% of East Asian population and in 0.3% of European population in the Genome Aggregation Database (gnomAD), whereas Arg1201Trp is present in 18.6% of the European population and only 0.6% in the Asian population. A high prevalence of double mutants is not observed except in Finnish population, where both mutants are present independently in 10.9% and 23.9% of the population (Karczewski *et al*., 2020).

The two SNPs (Arg996Gln and Arg1201Trp) can be involved in the closed-open conformation equilibrium and the described non-covalent oligomerization. However, no wide-ranging conformation differences were observed in the dimeric D3 mutants (Figure 4B), although minor alterations could be overlooked due to the low resolution in the flexible open conformation. Anyhow, the two mutations are likely to affect the molecular dynamics rather than the structure.

The mutation Arg996Gln, frequent in Asian population, affects the distance and the dynamics between the TIL3 domain and the VWD3, directly interfering in the conformational change and oligomerization. The open conformation was not detected in the D’D3CysD1 Arg99Gln cryoEM preparation during standard particle picking. Only template picking based on the WT open structure revealed a few particles. This observation, together with the described reduction of the distance between VWD3 and TIL3 domains, suggest that this mutation stabilizes the closed conformation and reduces the steric repulsion between Arg996 and TIL3. However, the VWD3 Arg996Gln mutant showed a higher occurrence of high order oligomers with an open conformation (Figure 4C), supporting that the opening conformation can still occur. When it formes, the absence of the positive charge residue near the newly exposed hydrophobic interaction face can modulate its properties. On the other hand, the Arg996Gln mutation produces a dramatic conformational change in the standard closed conformation tetramer. The His-tagged Arg996Gln mutant showed a high proportion of octameric particles in cryoEM that were essential for solving the high-resolution structure of the non-covalent interaction. The TIL3-TIL3 interaction is flexible in all cases, but the Arg99Gln mutation could add more freedom to allow the octameric arrangement. This mutation probably also increases the interaction affinity and therefore stabilizes the TIL interface, consequently also the second TIL3 domain in each disulfide-bonded dimer (Figure 4D) appears involved in non-covalent interactions making the formation of octamers possible. We could not find any significant disease association of the Arg99Gln mutation as the mutation has a low frequency in the used database (UK-Biobank). Further analysis on Arg996Gln and its potential association with disease is required.

The Arg1201Trp SNP, frequent in European population, shows a double role in the conformation alteration of the VWD3 assembly. In the closed conformation it establishes a novel interaction between the TIL3 domain and VWD3, and in the open conformation removes the possibility of the predicted salt bridge between Arg1201 and Glu981. By that we can speculate that this mutation should favor the closed conformation, even though open conformation molecules were still observed in cryoEM. The role of Arg1201Trp in MUC5AC VWD3 tetramerization is not fully understood. Arg1201 is located in the interaction site but not directly involved in any contact. However, it is in close proximity to Asp1195 and the limitations in resolution and flexibility of the interaction make it not possible to disregard the importance of the 1201 amino acid in the formation of the interface. On the other hand, the presence of the tryptophan interacting with VWD3 could affect the TIL3 flexibility that is important for the interaction. The double mutant shows that once Arg996 is mutated, Trp1201 stabilizes the interface bringing TIL3 closer to VWD3. In theory Arg1201Trp should limit the TIL3 flexibility and have the opposite effect of Arg996Gln. Nevertheless, we found a significant association between the Arg1201Trp SNP and COPD supporting that the mutation could favor tetramerization and thus net-like formation. This mutation has also recently been found to be associated with dysregulated inflammatory responses across keratoconic cone where MUC5AC mucin plays a protective role (Jaskiewicz *et al*., 2023). Arg996Gln and Arg1201Trp have thus the same effects in the closed-open conformation equilibrium, stabilizing the closed form. In the same way, the effect of both mutations in tetramerization seems to be equivalent even if the outcome has not been completely elucidated.

That the Arg1201Trp SNP promotes tetramerization suggests that it should increase mucus crosslinking. The amount of MUC5AC is known to be increased at lung diseases and especially COPD (Fahy *et al*., 2010;Fernandez-Blanco *et al*., 2018;Radicioni *et al*., 2021) and its role in trapping bacteria could be increased (Ermund *et al*., 2021). MUC5AC is known to anchor the MUC5B bundled strands to the surface goblet cells and a more cross-linked MUC5AC might increase the attachment and retard clearance (Ermund *et al*., 2017). That there is an association of the Arg1201Trp SNP with COPD suggests a close association between the MUC5AC mucin properties and the risk COPD.

The strong association of the MUC5B promotor SNP (rs35705950) leading to increased levels of MUC5B protein and IPF has been known for some time (Moll *et al*., 2023;Seibold *et al*., 2011). We here observed a strong association between the MUC5AC Arg1201Trp SNP and IPF. The fact that this has been overlooked might at first be a surprise, but this is explained by poor sequence coverage close to the repetitive and polymorphic parts of the *MUC5AC* gene. As the MUC5B and MUC5AC SNPs are located close together on the same chromosome and show linkage disequilibrium it is not a surprise that the MUC5AC SNP seems to have an association to IPF. However, interestingly it is only when both SNPs appear at the same time that they show a strong association with IPF. It is thus likely that the strong association of the MUC5B SNP (rs35705950) with IPF is caused by combination with the MUC5AC SNP (rs878913005).

In the normal lung, the small airways without submucosal glands are characterized by surface goblet cells producing only MUC5B mucin (Hill *et al*., 2022;Okuda *et al*., 2021). However, recent observations suggest that the MUC5AC mucin is increased in peripheral airways in IPF (Herrera *et al*., 2023). The reason for the association of the MUC5B promotor SNP with IPF has been suggested to be due to increased levels of the MUC5B mucin (Hancock *et al*., 2018). Although this is a likely link to IPF, an association of IPF susceptibility with a more cross-linked MUC5AC is even easier to understand. A more cross-linked MUC5AC might be more difficult to remove from the thin diameter peripheral airways and might lead to mucus retardation and with this a likely increased bacterial retardation. Previous observations that show the importance of MUC5B for IPF has been puzzling as the MUC5B promoter polymorphism was not associated with IPF in Asian population (Peljto *et al*., 2015). However, this is easier to explain by the present observation of the linkage of the two SNPs and the very low abundance of the MUC5AC Arg1201Trp SNP in the Asian population. This further supports the conclusion that it is the combination of the MUC5AC and MUC5B SNPs that is the required driver for IPF susceptibility.

## METHODS

### Production and purification of MUC5AC plasmids

The recombinant MUC5AC-D’-D3-CysD (WT and Arg996Gln), MUC5AC-D3-CysD (WT) and MUC5AC-D3 (WT and Arg996Gln) (GenBank accession number NM_001304359, residues 800-1481, 901-1481, 901-1366) plasmids were expressed with an N-terminal Hisx6 tag and a C-terminal Myc tag using the mammalian episomal expression vector pCEP-His. MUC5AC-D3 (WT, Arg996Gln, Arg1201Trp and Arg996Gln Arg1201Trp) was also expressed without any tag using the same expression vector.

CHO-Lec 3.2.8.1-S (Nilsson *et al*., 2014) cells were grown in 300 ml Freestyle™ CHO with 8 mM L-glutamine in an Erlenmeyer flask in 5% CO_2_. Transfection with NovaCHOice transfection kit (Merck, Nottingham, UK) was performed according to the manufacturer’s instructions. Four hours after transfection the temperature was decreased to 31°C. The supernatant was harvested after 48h by centrifugation for 10 min at 200 × g at room temperature and then immediately dialyzed against PBS 10 mM imidazole (His-tagged proteins) or 20 mM Tris pH 8 (His-tag free proteins) at 4°C. Several MUC5AC-D3 batches of 300 ml each were made in CHO-Lec 3.2.8.1-S.

MUC5AC was filtered (Durapore® Membrane Filter, 0.22 μm GVWP, Millipore) and further purified using an ÄKTA purifier (GE Healthcare). The His-tag containing proteins were loaded onto a HiTrap chelating HP nickel affinity 1-ml column (GE Healthcare). The bound components were eluted with a gradient of 10–300 mM imidazole in 20 mM Tris pH 7.4 and 150 mM NaCl. The His-tag free MUC5AC-D3 variants were loaded onto a HiPrep Q HP 16/10 anion exchange chromatography column (GE Healthcare) and eluted in a 0-500 mM NaCl gradient in 20 mM Tris pH 8.

The protein containing fractions were dialyzed against 20 mM Tris (pH 7.4) and 50 mM NaCl, loaded onto a Mono Q™ HR 10/10 anion exchange column (GE Healthcare) and eluted in a linear gradient from 50 to 500 mM NaCl. It was followed by size exclusion fractionation on a Superose 6 10/300 column (GE Healthcare) eluted in 20 mM Tris pH 7.4, 50 mM NaCl and 10 mM CaCl_2_ and collected in fractions of 0.5 ml.

Protein purity was checked by SDS-PAGE using 4-15% Mini-PROTEAN® TGX™ precast protein gels (Bio-Rad). Proteins were diluted in 2x SDS-PAGE loading buffer reaching a final concentration of 100 mM DTT or in DTT-free buffer, heated 5 minutes at 95 C and loaded into the gel together with the Precision Plus Protein Unstained Standard (Bio-Rad). Gels were stained with Coomassie brilliant blue G-250.

### Single particle cryoEM

In order to optimize the sample homogeneity only the central fraction of the size exclusion was used for all recombinant proteins sample preparation (MUC5AC-D’-D3-CysD1 (WT and Arg996Gln), MUC5AC-6xHis-D3 (WT and Arg996Gln) and MUC5AC-D3 (Arg1201Trp and Arg996Gln-Arg1201Trp). The protein concentration was adjusted to 0.6 µM in 20 mM Tris pH 7.4, 150 mM NaCl and 10 mM CaCl_2_.

The samples were loaded onto UltrAuFoil R1.2/1.3 300# (SPT Labtech) holey gold grids (D3 Arg1201Trp, D3 Arg996Gln-Arg1201Trp, 6xHis-D3 Arg996Gln and 6xHis-D3 WT) or Quantifoil Cu 1.2/1.3 300# (SPT Labtech) copper grids (D’-D3-CysD1 WT and D’-D3-CysD1 Arg996Gln) that were previously glow discharged at 15 mA for 40 s with a negative charge. The grids were plunge frozen using a Vitrobot Mark IV (ThermoFisher) set at 100% humidity and 4°C. 6xHis-D3 Arg996Gln data collections were performed in a Titan Krios microscope (Thermo Fisher) at 0.83 Å/pix and 300 kV acceleration voltage using a K2 Summit 4k x 4k detector (Gatan). 40 frames per movie were collected with an average electron dose per image of 1.15 e/Å^2^ and nominal defocus between -0.5 and -3.5 μm in 0.3 steps. The data from five grids collected using the same conditions were merged for reconstruction. The datasets from the other five recombinant proteins were collected using EPU software (Thermo Fisher Scientific) in Aberration-free image shift (AFIS) mode at 0.86 Å/pix and 300 kV acceleration voltage and a K3 Summit 6k x 4k detector (Gatan). 40 frames per movie were collected with an average electron dose per image of 1.25 (6xHis-D3 WT), 1.26 (D3 Arg996Gln Arg1201Trp), 1.29 (D’-D3-CysD1 WT and D’-D3-CysD1 Arg996Gln) or 1.49 e/Å^2^ (D3 Arg1201Trp) and defocus between -0.5 and -2.5 μm (D’-D3-CysD1 WT, 6xHis-D3 WT and D3 Arg996Gln Arg1201Trp) or -0.5 and -3.0 μm (D’-D3-CysD1 Arg996Gln and D3 Arg1201Trp). These parameters are sumarized in Supplementary Table S1. The micrographs were imported in cryoSPARC v.4.2.1 (Punjani *et al*., 2017) and patch motion corrected. CTF correction was performed in cryoSPARC using Patch CTF. The micrographs were manually curated and outliers were removed. The data processing steps leading to the PDB deposited models are schematically represented in Supplementary Figures S1 to S5.

#### D’-D3-CysD1 WT

A first round of manual picking was performed. The particles were 2D classified and the bests classes were used for template-based automatic particle picking. The particles were extracted using a 256 pixel box size and 2D classified. The junk particles were removed and the rest were used for *ab-initio* reconstruction (2 classes) and heterogeneous refinement. A fraction of the particles from the best locking class was used for Topaz particle picking training (Bepler *et al*., 2019). The Topaz picking model was applied to the curated dataset and the particles were extracted using a 256 pixel box size and 2D classified for junk particle removal. The particles were used for *ab-initio* reconstruction (4 classes) and heterogeneous refinement. A fraction of the particles from the best locking class was used for a new Topaz particle picking training. A new particle extraction was performed and the previous steps were repeated generating new 4 *ab-initio* classes that were used again in heterogeneous refinement. The heterogeneous refinement classes were used as templates for non-uniform refinement using C1 and C2 symmetries. The particles from the highest resolution class in the non-uniform refinement with C2 symmetry were further 3D classified by a new *ab-initio* reconstruction round in 2 classes and heterogeneous refinement. The best quality class was non-uniform refined imposing C2 symmetry that was used as an input for a final local refinement also imposing C2 symmetry.

#### D’-D3-CysD1 Arg996Gln

The same protocol as for D’-D3-CysD1 WT Dimers was applied until the first Topaz particle picking and extraction. Then the particles were directly classified in 2 *ab-initio* classes. The particles from the best class were 2D classified and only the high-resolution classes were selected. These particles were used for a new Topaz training and the process was repeated. The particles were used in a new *ab-initio* reconstruction round in 2 classes and heterogeneous refinement. The best quality class was non-uniformly refined imposing C2 symmetry that was used as an input for a global CTF refinement and a final local refinement also imposing C2 symmetry. To look for the presence of open dimers in the sample, D’-D3-CysD1 WT open dimer templates were used for template-based automatic particle picking. Particles were extracted using a 256 pixel box size and 2D classified for junk particle removal. The remaining particles were 3D classified by *ab-initio* reconstruction in 2 classes and heterogeneous refinement, and further 2D classified.

#### D3 Arg1201Trp

An initial blob picking was performed using 1,000 micrographs setting a minimum particle diameter of 70 Å and maximum of 140 Å. The particles were 2D classified and the bests classes were used for template-based automatic particle picking. The particles were extracted using a 256 pixel box size and 2D classified. The junk particles were removed and the rest were used for *ab-initio* reconstruction (2 classes) and heterogeneous refinement. A fraction of the particles from the best locking class was used for Topaz particle picking training. The Topaz picking model was applied to the curated dataset and the particles were extracted using 256 pixels box size and 2D classified for junk particle removal. Particles were divided in two *ab-initio* classes. A fraction of the particles from the best locking class was used for a new Topaz particle picking training. A new particle extraction was performed and the previous steps were repeated generating new 2 *ab-initio* classes that were used in heterogeneous refinement. The particles from the highest quality class were 3D classified again by *ab-initio* reconstruction in two classes and heterogeneous refinement. The best class was non-uniform refined imposing C2 symmetry that was used as an input for a global CTF refinement and a final local refinement also imposing C2 symmetry.

#### D3 Arg996Gln Arg1201Trp

The same protocol than in D’-D3-CysD1 WT Dimers was applied changing the *ab-initio* and heterogeneous refinement with 4 classes for 2 classes and skipping the further repetition.

#### 6xHis-D3 Arg996Gln

The same protocol as for D’-D3-CysD1 R996 Dimers, but using a 450 pixel box size, was applied until the *ab-initio* reconstruction following the first Topaz particle picking and extraction step. Both *ab-initio* classes, one representing the tetrameric conformation and the other the octameric, were further 2D classified to remove junk particles and used independently in a second Topaz training. Both models were used for particle picking and two group of particles were newly extracted. The particles based in the tetrameric conformation training were 3D *ab-initio* reconstructed in 2 classes and further heterogeneously refined. The particles from the heterogeneous refinement class corresponding to a tetramer were 3D *ab-initio* reconstructed in 2 classes and further heterogeneously refined again. The best quality class was non-uniformly refined. Finally, the non-uniform refinement was used as an input for a final local refinement. In parallel, the particles based in the octameric conformation training were also 3D *ab-initio* reconstructed in 2 classes and further heterogeneously refined. The particles from the heterogeneous refinement class corresponding to an octamer were 3D *ab-initio* reconstructed in 2 classes and further heterogeneously refined again. The best quality class was non-uniformly refined. The 3D volume was used to create two masks in Chimera (Pettersen *et al*., 2004), one including the best defined tetramer in the octameric assembly (A) and the inverted mask (B). The masks ware newly imported in cryoSPARC and the mask A was dilated 6 Å and soft padded 8 Å. The particles from the non-uniform refinement were 3D classified in 4 classes using the non-uniformly refinement mask as a solvent mask and the mask A as focused mask. All four classes were further non-uniform refined. The particles and volume from the best resolution class were used for particle subtraction applying mask B. The newly generated particles together with the non-uniform refinement volume and the mask A were used as inputs in a last local refinement step.

#### D’-D3-CysD1 WT, D’-D3-CysD1 Arg996Gln, D3 Arg1201Trp, D3 Arg996Gln and 6xHis-D3 WT Tetramers

Manual and template picking was performed as for most of the dimers but using a bigger box size, 320, 450 or 512 pixels. Particles were 2D classified and the best classes were used for Topaz particle picking. Particles were extracted using the same box size and further 2D classified.

### Model building

An initial model of MUC5AC-D3 Arg996Gln dimer was built in AlphaFold2 (Jumper *et al*., 2021). It was fitted into the density map with Molrep (Vagin and Teplyakov, 1997) and manually built using Coot 0.9.8.1. The model was refined along different iterations using the real-space refinement tool in Phenix 1.20.1-4487 (Liebschner *et al*., 2019) and manual refinement in Coot. The final structure refinement validation was performed by Molprobity (Williams *et al*., 2018). The other models were built following the same protocol but using the final MUC5AC-D3 Arg996Gln dimer model (WT and Arg996Gln Arg1201Trp dimers, and Arg996Gln tetramer) or the WT dimer model (Arg1201Trp) as initial models. PyMol (Schrodinger, LLC (2015) The PyMOL Molecular Graphics System, version 2.5) and UCSF ChimeraX (Meng *et al*., 2023) were used for structure analysis and figures generation.

The MUC5AC-D3 open conformation model was generated manually fitting the MUC5AC-D3 WT closed conformation C8-3 and TIL3 dimer and two independent VWD3 domains into the low-resolution density assisted by the fit in map tool. The model building was finished using ISOLDE (Croll, 2018).

### Human samples

Stomach biopsies were acquired during routine endoscopies at the Sahlgrenska University Hospital (ethical permission 085-06). All subjects gave written informed consent. Samples were fixed directly in Carnoýs solution (composition 60% methanol, 30% chloroform and 10% glacial acetic acid). Lung samples were acquired during resection of diseased lungs at transplantation (ethical permission 2020-03693). Pieces of lung tissue were fixed in 10% neutral buffered formalin according to routine procedures at the Sahlgrenska University Hospital. Fixed human tissue was transferred to 70% ethanol and stored until paraffin embedding, sectioning into 4 µm sections and mounting on superfrost plus slides (Cat# 631-9483, VWR, Avantor, Radnor Township, PA). Researchers had access to medical records.

### Immunofluorescent staining of histological sections

Sections were baked on the slides at 60°C for 2 hours, dewaxed using xylene and hydrated. Antigen heat-induced epitope retrieval was performed with 10 mM citrate buffer pH 6.0 at 100°C for 20 min and then room temperature for another 20 min. Sections were washed in PBS, a barrier was drawn with an ImmEdge® Hydrophobic Barrier PAP Pen (H-4000, Vector laboratories, Newark, CA). Unspecific epitopes were blocked with 3% donkey serum in Tris-buffered saline and sections were permeabilized with 0.1% Triton X-100. Stainings were performed with sequential incubation with custom made polyclonal rabbit anti-human MUC5B antibodies (1:200) in block solution (Fakih *et al*., 2020) overnight at 4°C and monoclonal mouse anti-human MUC5AC (Lidell *et al*., 2008)(1:200) in block solution over night at 4°C (Cat# ab3649, Abcam, Cambridge, UK, RRID:AB_2146844). Secondary antibody was polyclonal donkey anti-rabbit Alexa Fluor 488 (Cat# A-21206, Thermo Fisher Scientific, Waltham, MA, RRID:AB_2535792) and polyclonal donkey anti-mouse Alexa Fluor 647 (Cat# A-31571, Thermo Fisher Scientific, RRID:AB_162542). Nuclei were staned with Hechts 34580 (Thermo Fisher Scientific)

### Imaging

Images were acquired using ZEISS ZEN Microscopy Software (Carl Zeiss, Oberkochen, Germany, RRID:SCR_013672) on the ZEISS LSM900 with Airyscan 2 (Carl Zeiss, Oberkochen, Germany, RRID:SCR_022263) and processed with Imaris version software (Oxford Instruments, Abingdon, UK, RRID:SCR_007370).

### Piglet airway tissue

Ethical permitssion for experiments involving newborn piglets (Sus scrofa domesticus) were obtained from Regierungen von Oberbayern, Munich, Germany (AZ55.2-1-54-2531-78-07) and Jordbruksverket, Jönköping, Sweden (Dnr 6.7.18-12708/2019). Tracheas were acquired from wild type piglets (Sus scrofa domesticus). To induce birth, intramuscular administration of 0.175 mg Cloprostenol (Estrumate®, Intervet GmbH, Unterschleissheim, Germany), on gestation day 112–114. Within 24 h of birth, piglets were anesthetized by Ketamine (Ursotamin®, Serumwerk Bernburg, Germany) and Azaperone (Stresnil®, Elanco Animal Health, Bad Homburg, Germany) and killed by intracardial injection of Tanax® T61 euthanasia solution (Intervet GmbH, Unterschleissheim, Germany). Tracheas from the larynx and lungs were excised and the lung parenchyma removed under Perfadex® solution, pH 7.2 (XVIVO Perfusion, Gothenburg, Sweden) before the prepared airways including larynx, trachea and bronchi were transferred to a 50 ml tube with Perfadex® solution, pH 7.2 and shipped at 4°C overnight to Gothenburg.

### Electron microscopy

Distal tracheal tissue (two to three cartilage rings in length) from newborn piglets were fixed in modified Karnovsky’s fixative (2% paraformaldehyde, 2.5% glutaraldehyde in 0.05 M sodium cacodylate buffer, pH 7.2) for 24 h at 4°C. Postfixation was performed in 1% OsO4 at 4 C three times with intervening 1% thiocarbohydrazide steps. The samples were dehydrated with increasing concentrations of ethanol followed by hexamethyldisilazane that was allowed to evaporate. Samples were mounted on aluminum specimen pin stubs (Cat# AGG301, Agar Scientific, Stansted, Essex, UK) with carbon tabs (Cat# AGG3347N, Agar Scientific, Stansted, Essex, UK) and conductive silver paint (Cat# 16040-30, Ted Pella, Redding, CA). To decrease charging, samples were sputter-coated with palladium before imaging at 3 kV in a field emission scanning electron microscope (Zeiss DSM 982 Gemini, Carl Zeiss, Oberkochen, Germany).

### UK-biobank

Two cohorts were defined using the UK-biobank Research Analysis Platform (DNAnexus), patients diagnosed with COPD (n=3,261) and patients diagnosed with IPF (n=107). A randomized control group (n=10,000) was stabilized with the rest of patients not included in these cohorts. The genetic information regarding chromosome 11 contained in the available variant call files (VCFs) was extracted (COPD n=1,139, IPF n=107 and control n=5,196). Annotations in GRCh38 position 1177533 (rs36189285, MUC5AC Arg996Gln), 1180138 (rs878913005, MUC5AC Arg1201Trp) and 1219991 (rs35705950, MUC5B promotor) were searched and participants were classified as SNP carriers (heterozygous and homozygous) or WT (wild type or not annotated). The database used was from August 29, 2022. The diseases association statistical analysis was performed using Fisher’s exact test.

## Data availability

The structural data from cryoEM is deposited at PDB and EMDB with accession 8QTV and 18654 (D’-D3-CysD1 WT), 8QTB and 18648 (D’-D3-CysD1 Arg996Gln), 8R1U and 18828 (D3 Arg1201Trp), 8R1Z and 18829 (D3 Arg996Gln Arg1201Trp) and 8QSP and 18638 (6xHis-D3 Arg996Gln).

## SUPPLEMENTAL INFORMATION

Supplemental information with supplemental Figures is included.

## ACKNOWLEDGEMENTS

This work was supported by the The Knut and Alice Wallenberg Foundation (2017.0028), European Research Council ERC (101100663, 694181), Swedish Research Council (2017-00958, 2022-00646), IngaBritt and Arne Lundberg Foundation (2018-0117), Sahlgren’s University Hospital (ALFGBG-440741), Bill and Melinda Gates Foundation (OPP1202459), The Swedish Heart-Lung Foundation (20220404 (AE); 20230413 (GCH)), and Wilhelm and Martina Lundgren’s Foundation. The data was collected at the cryoEM Swedish National Facility funded by the Knut and Alice Wallenberg, Family Erling Persson and Kempe Foundations, SciLifeLab, Stockholm University and Umeå University.

## AUTHOR CONTRIBUTIONS

ST-M, GCH initiated and planned research; STM designed, performed and analyzed structural experiments and results; AE performed tissue stainings and microscopy; JMM collected human samples; GCH obtained funding; ST-M and GCH wrote the manuscript with input from all authors that also approved the final text.

## REFERENCES

Asker N, Axelsson MAB, Olofsson SO, and Hansson GC (1998) Human MUC5AC mucin dimerizes in the rough endoplasmic reticulum, similarly to the MUC2 mucin. Biochem J, 335, 381–387.

Axelsson MAB, Asker N, and Hansson GC (1998) O-glycosylated MUC2 monomer and dimer from LS 174T cells are water-soluble, whereas larger MUC2 species formed early during biosynthesis are insoluble and contain nonreducible intermolecular bonds. J Biol Chem, 273, 18864–18870.

Bepler T, Morin A, Rapp M, Brasch J, Shapiro L, Noble AJ, and Berger B (2019) Positive-unlabeled convolutional neural networks for particle picking in cryo-electron micrographs. Nat Meth, 16, 1153–1160.

Bhattacharyya R, Dhar J, Dastidar SG, Chakrabartia P, and Weiss MS (2020) The susceptibility of disulfide bonds towards radiation damage may be explained by S-O interactions. IUCrJ, 7, 825–834.

Bos MF, Ermund A, Hansson GC, and de Graaf J (2023) Goblet-cell interactions reorient bundled mucus strands for efficient airway clearance. PNAS Nexus, 2, pgad388.

Carpenter J, Wang Y, Gupta R, Li Y, Haridass P, Subramani DB, Reidel B, Morton L, Ridley C, ONeal WK, Buisine MP, Ehre C, Thornton DJ, and Kesimer M (2021) Assembly and organization of the N-terminal region of mucin MUC5AC: Indications for structural and functional distinction from MUC5B. PNAS, 118, e2104490118.

Croll T (2018) ISOLDE: a physically realistic environment for model building into low-resolution electron-density maps. Acta Cryst D, 74, 519–530.

Dong X, Leksa NC, Chhabra ES, Arndt JW, Lu Q, Knockenhauer KE, Peters RT, and Springer TA (2019) The von Willebrand factor D’D3 assembly and structural principles for factor VIII binding and concatemer biogenesis. Blood, 133, 1523–1533.

Ermund A, Meiss LN, Dolan B, Jaudas F, Ewaldsson L, Bahr A, Klymiuk N, and Hansson GC (2021) Mucus threads from surface goblet cells clear particles from the airways. Respiratory Research, 22, 303.

Ermund A, Meiss LN, Dolan B, Bahr A, Klymiuk N, and Hansson GC (2018) The mucin bundles responsible for airway cleaning are retained in cystic fibrosis and by cholinergic stimulation. Eur Resp J, 52, 1800457.

Ermund A, Meiss LN, Rodriguez-Pineiro AM, Bahr A, Nilsson HE, Trillo-Muyo S, Ridley C, Thornton DJ, Wine JJ, Hebert H, Klymiuk N, and Hansson GC (2017) The normal trachea is cleaned by MUC5B mucin bundles from the submucosal glands coated with the MUC5AC mucin. Biochem Biophys Res Commun, 492, 331–337.

Fahy JV and Dickey BF (2010) Airway Mucus Function and Dysfunction. New England Journal of Medicine, 363, 2233–2247.

Fakih D, Rodriguez Pineiro AM, Trillo-Muyo S, Evans CM, Ermund A, and Hansson GC (2020) Normal murine respiratory tract has its mucus concentrated in clouds based on the Muc5b mucin. Am J Physiol Lung Cell Mol Physiol, 318, L1270–L1279.

Fernandez-Blanco JA, Fakih D, Arike L, Rodriguez-Pineiro A, Martinez-Abad B, Skansebo E, Jackson S, Root J, Singh D, McCrae C, Evans CM, Astrand A, Ermund A, and Hansson GC (2018) Attached stratified mucus separates bacteria from the epithelial cells in COPD lungs. J Clin Invest Insights, 3, e120994.

Gallego P, Garcia-Bonete MJ, Trillo-Muyo S, Recktenwald CV, Johansson MEV, and Hansson GC (2023) The intestinal MUC2 mucin C-terminus is stabilized by an extra disulfide bond in comparison to von Willebrand factor and other gel-forming mucins. Nature Communications, 14, 1969.

Hancock LA, Hennessy CE, Solomon GM, Dobrinskikh E, Estrella A, Hara N, Hill DB, Kissner WJ, Markovetz MR, Grove Villalon DE, Voss ME, Tearney GJ, Carroll KS, Shi Y, Schwarz MI, Thelin WR, Rowe SM, Yang IV, Evans CM, and Schwartz DA (2018) Muc5b overexpression causes mucociliary dysfunction and enhances lung fibrosis in mice. Nature Communications, 9, 5363.

Hansson GC (2020) Mucins and the Microbiome. Annu Rev Biochem, 89, 769–793.

Herrera JA, Dingle LA, Monetero MA, Venkateswaran RV, Blaikley JF, Granato F, Pearson S, Lawless C, and Thornton DJ (2023) Morphologically intact airways in lung fibrosis have an abnormal proteome. Respiratory Research, 24, 99.

Hill DB, Button B, Rubinstein M, and Boucher RC (2022) Physiology and Pathophysiology of Human Airway Mucus. Physiol Rev, 102, 1757--1836.

Hoang ON, Ermund A, Jaramillo AM, Fakih D, French CB, Flores JR, Karmouty-Quintana H, Magnusson JM, Fois G, Fauler M, Frick M, Braubach P, Hales JB, Kurten RC, Panettieri R, Vergara L, Ehre C, Adachi R, Tuvim MJ, Hansson GC, and Dickey BF (2022) Mucins MUC5AC and MUC5B Are Variably Packaged in the Same and in Separate Secretory Granules. Am J Resp Crit Care Med, 206, 1081–1095.

Hoegger MJ, Fischer AJ, McMenimen JD, Ostedgaard LS, Tucker AJ, Awadalla MA, Moninger TO, Michalski AS, Hoffman EA, Zabner J, Stoltz DA, and Welsh MJ (2014) Impaired mucus detachment disrupts mucociliary transport in a piglet model of cystic fibrosis. Science, 345, 818–822.

Jaskiewicz K, Maleszka-Kurpiel M, Kabza M, Karolak JA, and Gajecka M (2023) Sequence variants contributing to dysregulated inflammatory responses across keratoconic cone surface in adolescent patients with keratoconus. Front Immunol, 14, 1197054.

Javitt G, Calvo MLG, Albert L, Reznik N, Ilani T, Diskin R, and Fass D (2019) Intestinal Gel-Forming Mucins Polymerize by Disulfide-Mediated Dimerization of D3 Domains. J Mol Biol, 431, 3740–3752.

Javitt G, Khmelnitsky L, Albert L, Bigman LS, Elad N, Morgenstern D, Ilani T, Levy Y, Diskin R, and Fass D (2020) Assembly Mechanism of Mucin and von Willebrand Factor Polymers. Cell, 183, 717–729.

Jumper J, Evans R, Pritzel A, Green T, Figurnov M, Ronneberger O, Tunyasuvunakool K, Bates R, Zidek A, otapenko A, ridgland A, eyer C, ohl SAA, allard AJ, owie A, omera-Paredes B, ikolov S, ain R, dler J, ack T, etersen S, eiman D, lancy E, ielinski M, teinegger M, acholska M, erghammer T, odenstein S, ilver D, inyals O, enior AW, avukcuoglu K, ohli P, and assabis D (2021) Highly accurate protein structure prediction with AlphaFold. Nature, 596, 583–589.

Karczewski KJ, Francioli LC, Tiao G, Cummings BB, Alfoldi J, Wang Q, Collins RL, Laricchia KM, Ganna A, Birnbaum DP, Gauthier LD, Brand H, Solomonson M, Watts NA, Rhodes D, Singer-Berk M, England EM, Seaby EG, Kosmicki JA, Walters RK, Tashman K, Farjoun Y, Banks E, Poterba T, Wang A, Seed C, Whiffin N, Chong JX, Samocha KE, Pierce-Hoffman E, Zappala Z, ODonnell-Luria AH, Minikel EV, Weisburd B, Lek M, Ware JS, Vittal C, Armean IM, Bergelson L, Cibulskis K, Connolly KM, Covarrubias M, Donnelly S, Ferriera S, Gabriel S, Gentry J, Gupta N, Jeandet T, Kaplan D, Llanwarne C, Munshi R, Novod S, Petrillo N, Roazen D, Ruano-Rubio V, Saltzman A, Schleicher M, Soto J, Tibbetts K, Tolonen C, Wade G, Talkowski ME, Aguilar Salinas CA, Ahmad T, Albert CM, Ardissino D, Atzmon G, Barnard J, Beaugerie L, Benjamin EJ, Boehnke M, Bonnycastle LL, Bottinger EP, Bowden DW, Bown MJ, Chambers JC, Chan JC, Chasman D, Cho J, Chung MK, Cohen B, Correa A, Dabelea D, Daly MJ, Darbar D, Duggirala R, Dupuis J, Ellinor PT, Elosua R, Erdmann J, Esko T, Frkkil M, Florez J, Franke A, Getz G, Glaser B, Glatt SJ, Goldstein D, Gonzalez C, Groop L, Haiman C, Hanis C, Harms M, Hiltunen M, Holi MM, Hultman CM, Kallela M, Kaprio J, Kathiresan S, Kim BJ, Kim YJ, Kirov G, Kooner J, Koskinen S, Krumholz HM, Kugathasan S, Kwak SH, Laakso M, Lehtimeki T, Loos RJF, Lubitz SA, Ma RCW, MacArthur DG, Marrugat J, Mattila KM, McCarroll S, McCarthy MI, McGovern D, McPherson R, Meigs JB, Melander O, Metspalu A, Neale BM, Nilsson PM, ODonovan MC, Ongur D, Orozco L, Owen MJ, Palmer CNA, Palotie A, Park KS, Pato C, Pulver AE, Rahman N, Remes AM, Rioux JD, Ripatti S, Roden DM, Saleheen D, Salomaa V, Samani NJ, Scharf J, Schunkert H, Shoemaker MB, Sklar P, Soininen H, Sokol H, Spector T, Sullivan PF, Suvisaari J, Tai ES, Teo YY, Tiinamaija T, Tsuang M, Turner D, Tusie-Luna T, Vartiainen E, Vawter MP, Ware JS, Watkins H, Weersma RK, Wessman M, Wilson JG, Xavier RJ, Neale BM, Daly MJ, MacArthur DG, and Genome Aggregation Database Consortium (2020) The mutational constraint spectrum quantified from variation in 141,456 humans. Nature, 581, 434–443.

Kellokumpu S (2019) Golgi pH, Ion and Redox Homeostasis: How Much Do They Really Matter? Frontiers Cell Developmental Biology, 7, 93.

Lidell ME, Bara J, and Hansson GC (2008) Mapping of the 45M1 epitope to the C-terminal cysteine-rich part of the human MUC5AC mucin. FEBS J, 275, 481–489.

Liebschner D, Afonine PV, Baker ML, Bunkoczi G, Chen VB, Croll TI, Hintze B, Hung LW, Jain S, McCoy AJ, Moriarty NW, Oeffner RD, Poon BK, Prisant MG, Read RJ, Richardson JS, Richardson DC, Sammito MD, Sobolev OV, Stockwell DH, Terwilliger TC, Urzhumtsev AG, Videau LL, Williams CJ, and Adams PD (2019) Macromolecular structure determination using X-rays, neutrons and electrons: recent developments in Phenix. Acta Cryst D, 75, 861–877.

Meng EC, Goddard TD, Pettersen EF, Couch GS, Pearson ZJ, Morris JH, and Ferrin TE (2023) UCSF ChimeraX: Tools for structure building and analysis. Protein Science, 32, e4792.

Moll M, Peljto AL, Kim JS, Xu H, Debban CL, Chen X, Menon A, Putman RK, Ghosh AJ, Saferali A, Nishino M, Hatabu H, Hobbs BD, Hecker J, McDermott G, Sparks JA, Wain LV, Allen RJ, Tobin MD, Raby BA, Chun S, Silverman EK, Zamora AC, Ortega VE, Garcia CK, Barr RG, Bleecker ER, Meyers DA, Kaner RJ, Rich SS, Manichaikul A, Rotter JI, Dupuis J, OConnor GT, Fingerlin TE, Hunninghake GM, Schwartz DA, and Cho MH (2023) A Polygenic Risk Score for Idiopathic Pulmonary Fibrosis and Interstitial Lung Abnormalities. Am J Resp Crit Care Med, 208, 791–801.

Nilsson HE, Ambort D, Backstrom M, Thomsson E, Koeck PJ, Hansson GC, and Hebert H (2014) Intestinal MUC2 mucin supramolecular topology by packing and release resting on D3 domain assembly. J Mol Biol, 426, 2567–2579.

Okuda K, Chen G, Subramani DB, Wolf M, Gilmore RC, Kato T, Radicioni G, Kesimer M, Chua M, Dang H, Livraghi-Butrico A, Ehre C, Doerschuk CM, Randell SH, Matsui H, Nagase T, O’Neal WK, and Boucher RC (2018) Localization of Secretory Mucins MUC5AC and MUC5B in Normal/Healthy Human Airways. Am J Resp Crit Care Med, 199, 715–727.

Okuda K, Dang H, Kobayashi Y, Carraro G, Nakano S, Chen G, Kato T, Asakura T, Gilmore RC, Morton LC, Lee RE, Mascenik T, Yin WN, Barbosa Cardenas SM, ONeal YK, Minnick CE, Chua M, Quinney NL, Gentzsch M, Anderson CW, Ghio A, Matsui H, Nagase T, Ostrowski LE, GRUBB BR, Olsen JC, Randell SH, Stripp BR, Tata PR, O’Neal WK, and Boucher RC (2021) Secretory Cells Dominate Airway CFTR Expression and Function in Human Airway Superficial Epithelia. American Journal of Respiratory and Critical Care Medicine, 203, 1275–1289.

Peljto AL, Selman M, Kim DS, Murphy E, Tucker L, Pardo A, Lee JS, Ji W, Schwarz MI, Yang IV, Schwartz DA, and Fingerlin TE (2015) The MUC5B Promoter Polymorphism Is Associated With Idiopathic Pulmonary Fibrosis in a Mexican Cohort but Is Rare Among Asian Ancestries. Chest, 147, 460–464.

Pettersen EF, Goddard TD, Huang CC, Couch GS, Greenblatt DM, Meng EC, and Ferrin TE (2004) UCSF Chimera−A visualization system for exploratory research and analysis. J Comput Chem, 25, 1605–1612.

Punjani A, Rubinstein JL, Fleet DJ, and Brubaker MA (2017) cryoSPARC: algorithms for rapid unsupervised cryo-EM structure determination. Nat Meth, 14, 290–296.

Radicioni G, Ceppe A, Ford AA, Alexis NE, Barr RG, Bleecker ER, Christenson SA, Cooper CB, Han MK, Hansel NN, Hastie AT, Hoffman EA, Kanner RE, Martinez FJ, Ozkan E, Paine R, Woodruff PG, O’Neal WK, Boucher RC, and Kesimer M (2021) Airway mucin MUC5AC and MUC5B concentrations and the initiation and progression of chronic obstructive pulmonary disease: an analysis of the SPIROMICS cohort. Lancet Respir Med, 9, 1241–1254.

Recktenwald CV, Karlsson G, Garcia-Bonete MJ, Katona G, Jensen M, Lymer R, Backstrom M, Johansson MEV, Hansson GC, and Trillo-Muyo S (2024) The structure of the second CysD domain of MUC2 and role in mucin organization by transglutaminase-based cross-linking. Cell Reports, 43, 114207.

Ridley C, Kirkham S, Williamson SJ, Davis CW, Woodman P, and Thornton DJ (2016) Biosynthesis of the Polymeric Gel-Forming Mucin MUC5B. Am J Physiol Lung Cell Mol Physiol, 310, L993–L1002.

Roy MG, Livraghi-Butrico A, Fletcher AA, McElwee MM, Evans SE, Boerner RM, Alexander SN, Bellinghausen LK, Song AS, Petrova YM, Tuvim MJ, Adachi R, Romo I, Bordt AS, Bowden MG, Sisson JH, Woodruff PG, Thornton DJ, Rousseau K, De la Garza MM, Moghaddam SJ, Karmouty-Quintana H, Blackburn MR, Drouin SM, Davis CW, Terrell KA, GRUBB BR, ’Neal WK, Flores SC, Cota-Gomez A, Lozupone CA, Donnelly JM, Watson AM, Hennessy CE, Keith RC, Yang IV, Barthel L, Henson PM, Janssen WJ, Schwartz DA, Boucher RC, Dickey BF, and Evans CM (2014) Muc5b is required for airway defence. Nature, 505, 412–416.

Ryan A, Smith A, Moore P, McNally S, Carrington SD, Reid CJ, and Clyne M (2015) Expression and characterisation of a novel recombinant version of the secreted human mucin MUC5AC in airway cell lines. Biochemistry, 54, 1089–1099.

Sabo MC, Thuong NTT, Chang X, Ardiansyah E, Tram TTB, Hai HT, Nghia HDT, Bang ND, Dian S, Ganiem AR, Shaporifar S, Kumar V, Li Z, Hibberd M, Khor CC, Thwaites GE, Heemskerk D, van Laarhoven A, van Crevel R, Dunstan SJ, and Shah JA (2023) MUC5AC Genetic Variation Is Associated With Tuberculous Meningitis Cerebral Spinal Fluid Cytokine Responses and Mortality. The Journal of Infectious Diseases, 228, 343–352.

Sadler JE (1998) Biochemistry and genetics of von Willebrand factor. Ann Rev Biochem, 67, 395–424.

Schutte A, Ermund A, Becker-Pauly C, Johansson MEV, Rodriguez-Pineiro AM, Backhed F, Muller S, Lottaz D, Bond JS, and Hansson GC (2014) Microbial Induced Meprin beta Cleavage in MUC2 Mucin and Functional CFTR Channel are Required to Release Anchored Small Intestinal Mucus. Proc Natl Acad Sci USA, 111, 12396–12401.

Seibold MA, Wise AL, Speer MC, Steele MP, Brown KK, Loyd JE, Fingerlin TE, Zhang W, Gudmundsson G, Groshong SD, Evans CM, Garantziotis S, Adler KB, Dickey BF, du Bois RM, Yang IV, Herron A, Kervitsky D, Talbert JL, Markin C, Park J, Crews AL, Slifer SH, Auerbach S, Roy MG, Lin J, Hennessy CE, Schwarz MI, and Schwartz DA (2011) A Common MUC5B Promoter Polymorphism and Pulmonary Fibrosis. New England Journal of Medicine, 364, 1503–1512.

Shrine N, Portelli MA, John C, Soler Artigas Ma, Bennett N, Hall R, Lewis J, Henry AP, Billington CK, Ahmad A, Packer RJ, Shaw D, Pogson ZEK, Fogarty A, McKeever TM, Singapuri A, Heaney LG, Mansur AH, Chaudhuri R, Thomson NC, Holloway JW, Lockett GA, Howarth PH, Djukanovic R, Hankinson J, Niven R, Simpson A, Chung KF, Sterk PJ, Blakey JD, Adcock IM, Hu S, Guo Y, Obeidat M, Sin DD, van den Berge M, Nickle DC, Bosse Y, Tobin MD, Hall IP, Brightling CE, Wain LV, and Sayers I (2019) Moderate-to-severe asthma in individuals of European ancestry: a genome-wide association study. The Lancet Respiratory Medicine, 7, 20–34.

Svensson F, Lang T, Johansson MEV, and Hansson GC (2018) The central exons of the human MUC2 and MUC6 mucins are highly repetitive and variable in sequence between individuals. Scientific Reports, 8, 17503.

Vagin A and Teplyakov A (1997) MOLREP: an Automated Program for Molecular Replacement. J Appl Cryst, 30, 1022–1025.

Weik M, Ravelli RBG, Kryger G, McSweeney S, Raves ML, Harel M, Gros P, Silman I, Kroon J, and Sussman JL (2000) Specific chemical and structural damage to proteins produced by synchrotron radiation. PNAS, 97, 623–628.

Widdicombe JH and Wine JJ (2015) Airway Gland Structure and Function. Physiol Rev, 95, 1241–1319.

Williams CJ, Headd JJ, Moriarty NW, Prisant MG, Videau LL, Deis LN, Verma V, Keedy DA, Hintze BJ, Chen VB, Jain S, Lewis SM, Arendall III WB, Snoeyink J, Adams PD, Lovell SC, Richardson JS, and Richardson DC (2018) MolProbity: More and better reference data for improved all-atom structure validation. Protein Science, 27, 293–315.

Yeshaya N, Gupta PK, Dym O, Morgenstern D, Major DT, and Fass D (2024) VWD domain stabilization by autocatalytic Asp-Pro cleavage. Protein Science, 33, e4929.

