## Supplement for "Structural mechanism of MUC5AC mucin net-like polymer formation and its SNP variability that affect risk of the lung diseases COPD and IPF"

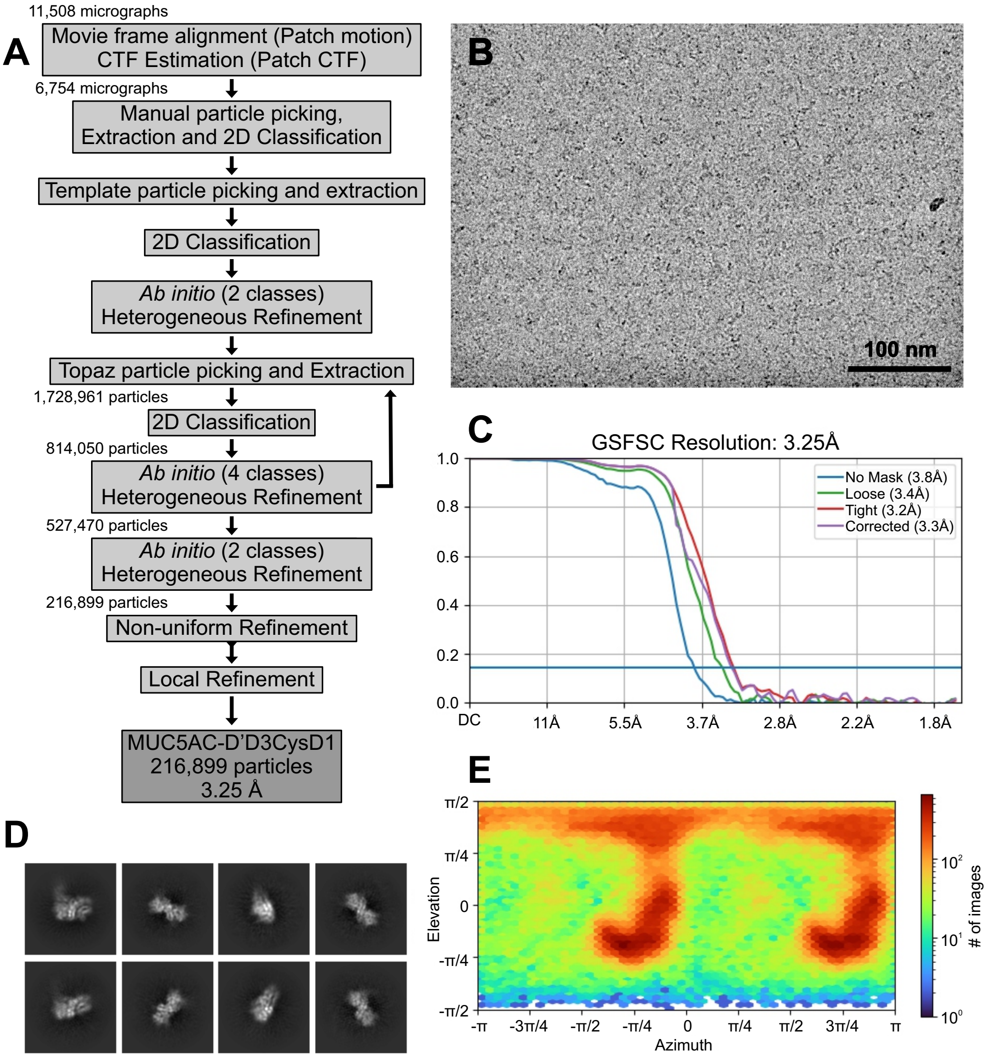


**Supplementary Figure 1. CryoEM structure of MUC5AC-D´D3CyD1.**

(A) Flowchart summary of cryoEM processing steps. The number of micrographs used and the number of particles remaining after each sorting step for the last iteration are specified.

(B) Representative micrograph. Scale bar is shown.

(C) Fourier shell correlations (FSC) for the final density map.

(D) Representative 2D classes.

(E) Per-particle distribution over azimuth and elevation angles for the final density map.

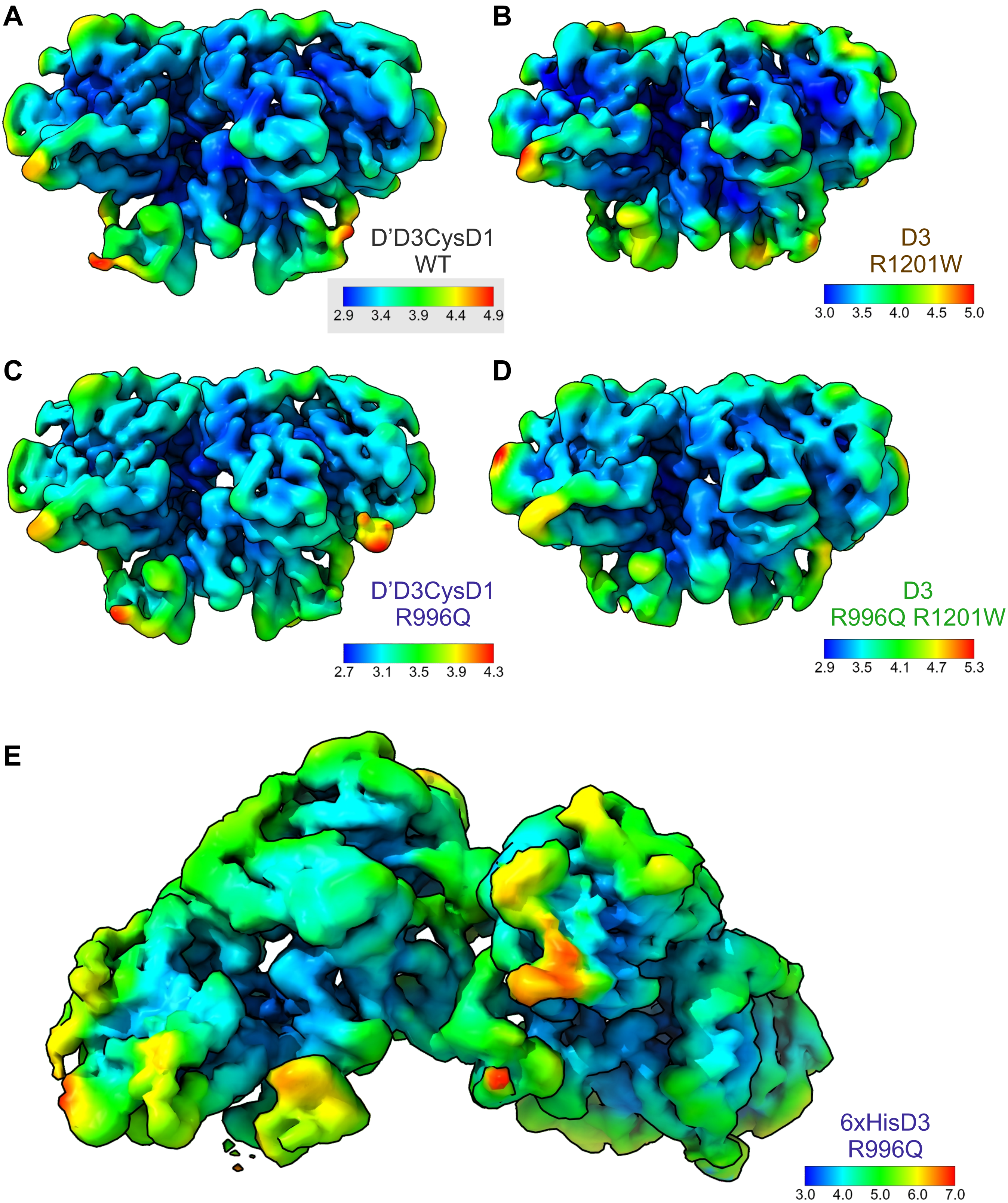


**Supplementary Figure 2. MUC5AC cryoEM maps of colored by local resolution.** The color keys show local resolution in Angstrom (Å). The keys were automatically generated showing the full range resolution calculated at FSC threshold 0.5.
(A) MUC5AC D'D3CysD1 WT.
(B) MUC5AC D3 R1201W.
(C) MUC5AC D'D3CysD1 R996Q.
(D) MUC5AC D3 R996Q R1201W.
(E) MUC5AC 6xHisD3 R996Q.

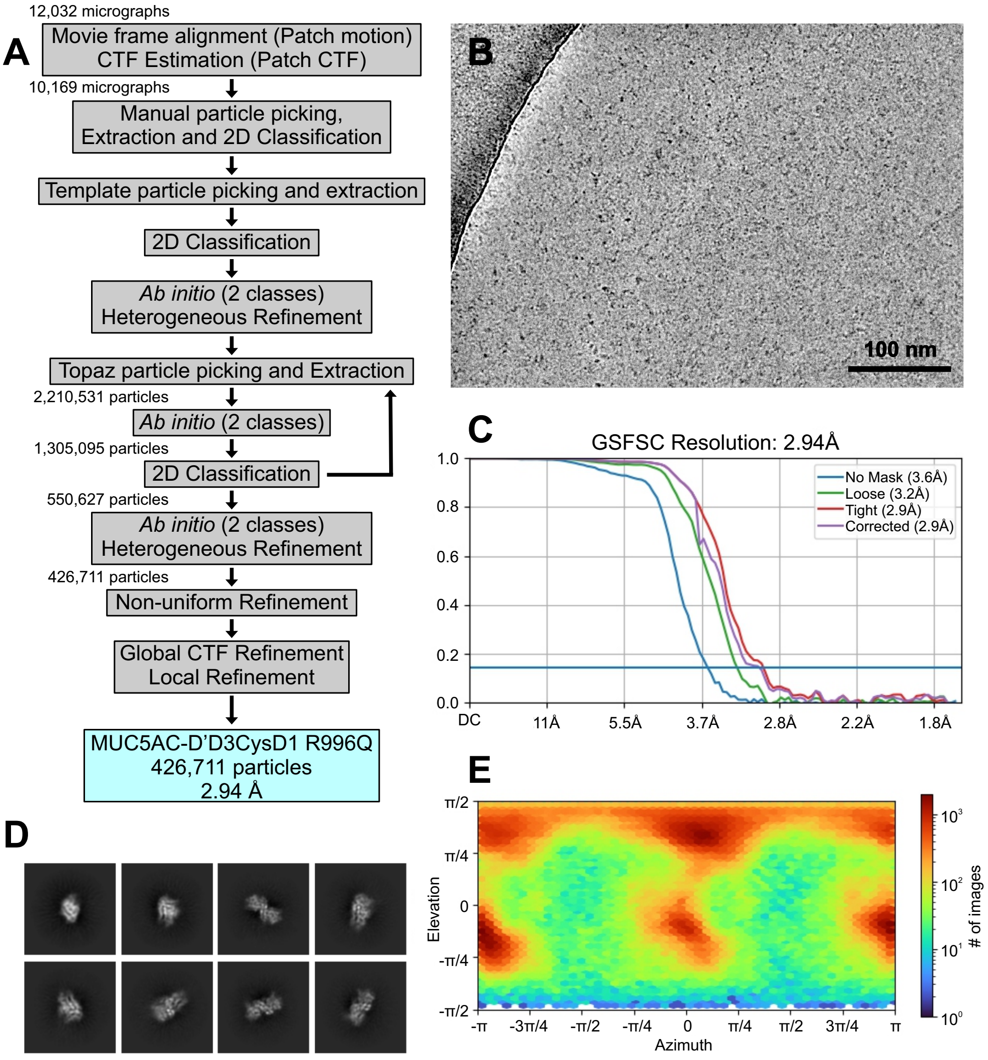


**Supplementary Figure 3. CryoEM structure of MUC5AC-D´D3CyD1 R996Q.**
(A) Flowchart summary of cryoEM processing steps. The number of micrographs used and the number of particles remaining after each sorting step for the last iteration are specified.
(B) Representative micrograph. Scale bar is shown.
(C) Fourier shell correlations (FSC) for the final density map.
(D) Representative 2D classes.
(E) Per-particle distribution over azimuth and elevation angles for the final density map.

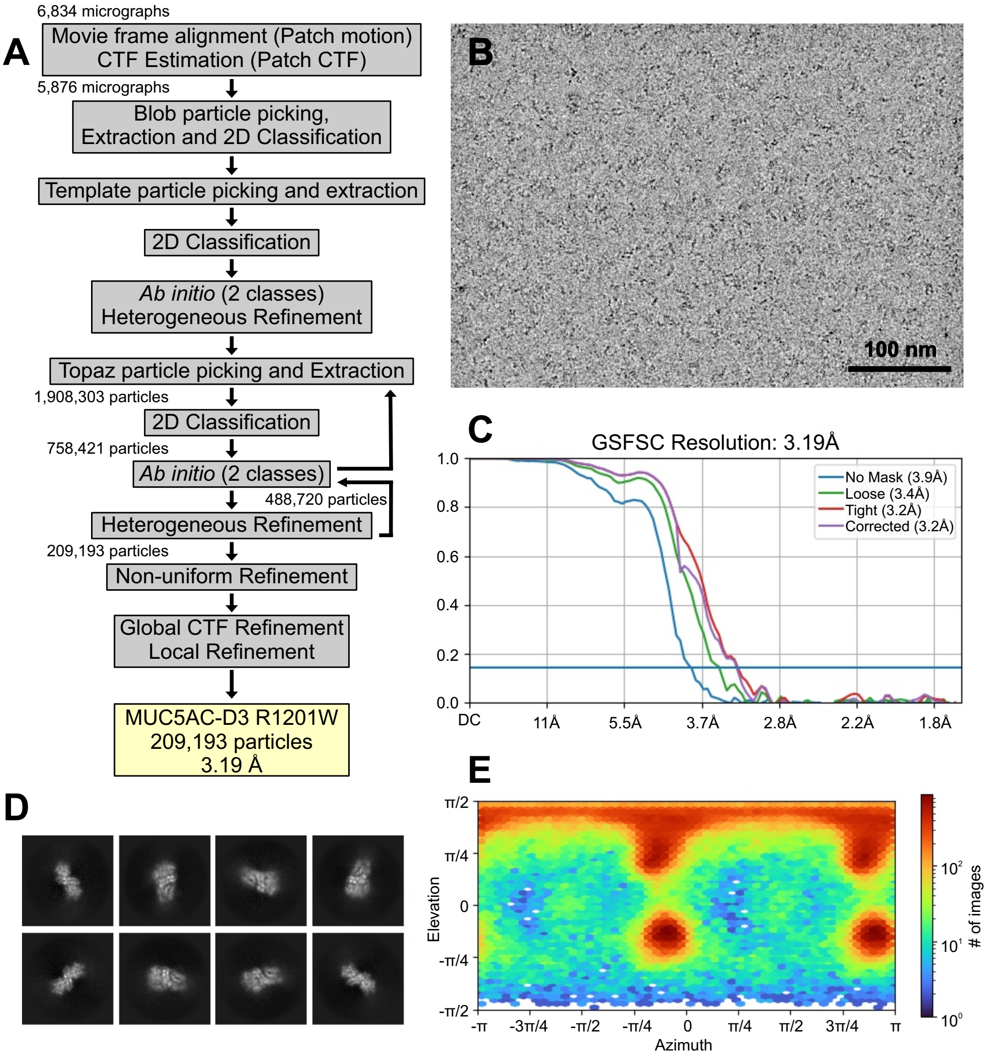


**Supplementary Figure 4. CryoEM structure of MUC5AC-D3 R1201W.**

(A) Flowchart summary of cryoEM processing steps. The number of micrographs used and the number of particles remaining after each sorting step for the last iteration are specified.

(B) Representative micrograph. Scale bar is shown.

(C) Fourier shell correlations (FSC) for the final density map.

(D) Representative 2D classes.

(E) Per-particle distribution over azimuth and elevation angles for the final density map.

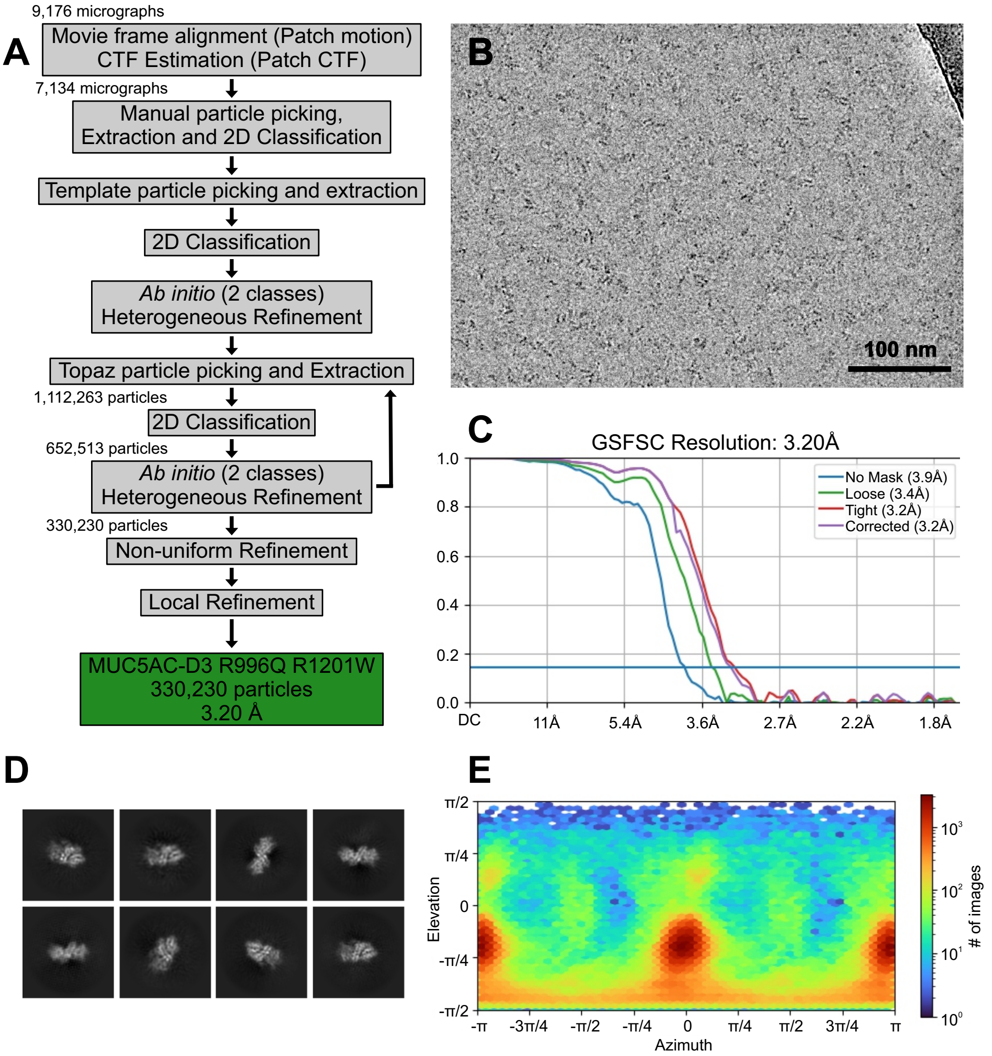


**Supplementary Figure 5. CryoEM structure of MUC5AC-D3 R996Q R1201W.**
(A) Flowchart summary of cryoEM processing steps. The number of micrographs used and the number of particles remaining after each sorting step for the last iteration are specified.
(B) Representative micrograph. Scale bar is shown.
(C) Fourier shell correlations (FSC) for the final density map.
(D) Representative 2D classes.
(E) Per-particle distribution over azimuth and elevation angles for the final density map.

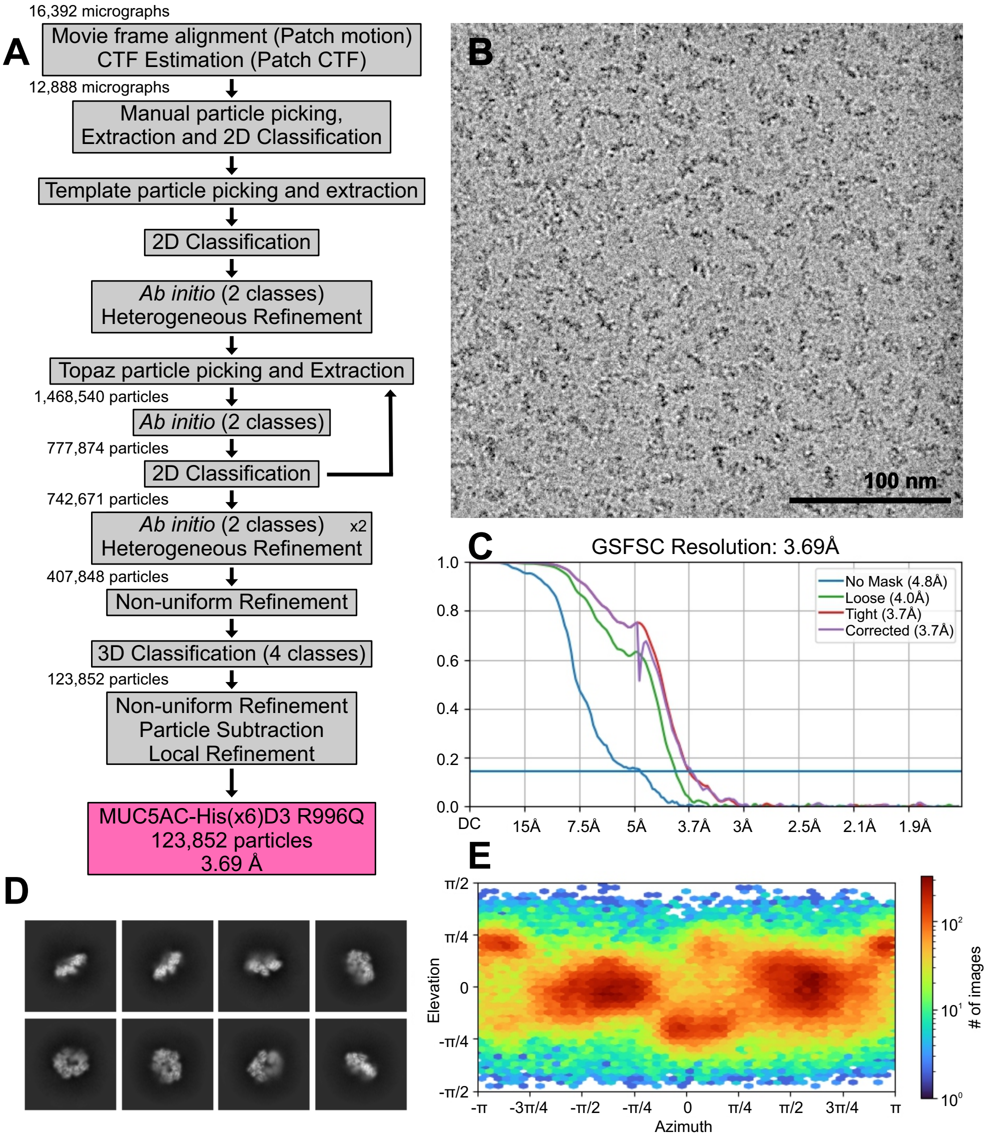


**Supplementary Figure 6. CryoEM structure of MUC5AC-6xHis-D3 R996Q.**
(A) Flowchart summary of cryoEM processing steps. The number of micrographs used and the number of particles remaining after each sorting step for the last iteration are specified.
(B) Representative micrograph. Scale bar is shown.
(C) Fourier shell correlations (FSC) for the final density map.
(D) Representative 2D classes.
(E) Per-particle distribution over azimuth and elevation angles for the final density map.

**Supplementary Table S1.** MUC5AC cryo-electron microscopy parameters.

| **Structure** | **D'D3CysD1 WT** | **D'D3CysD1 R996Q** | **D3 R1201W** | **D3 R996Q R1201W** | **His(x6)D3 R996Q** |
|---|---|---|---|---|---|
| **Data Accession** | | | | | |
| PDB | 8QTV | 8QTB | 8R1U | 8R1Z | 8QSP |
| EMDB | EMD-18654 | EMD-18648 | EMD-18828 | EMD-18829 | EMD-18638 |
| **Data Collection** | | | | | |
| Microscope | TFS KRIOS | TFS KRIOS | TFS KRIOS | TFS KRIOS | TFS KRIOS |
| Voltage (kV) | 300 | 300 | 300 | 300 | 300 |
| Detector | GATAN K3 | GATAN K3 | GATAN K3 | GATAN K3 | GATAN K2 |
| Pixel Size (Å) | 0.86 | 0.86 | 0.86 | 0.86 | 0.83 |
| Electron exposure ($e^-$/Å$^2$) | 50 | 51.6 | 50 | 50.3 | 46.2 |
| Defocus range (µm) | -0.5 to -2.5 | -0.5 to -3.0 | -0.5 to -3.0 | -0.5 to -2.5 | -0.5 to -3.5 |
| Micrographs | 11,508 | 12,032 | 6,834 | 9,176 | 16,392 |
| **Reconstruction** | | | | | |
| Software | CryoSPARC (v3.2) | CryoSPARC (v3.2) | CryoSPARC (v3.2) | CryoSPARC (v3.2) | CryoSPARC (v3.2) |
| Micrographs used | 6,754 | 10,169 | 5,876 | 7,134 | 12,888 |
| Particles used in refinement | 216,899 | 426,711 | 209,193 | 330,230 | 123,852 |
| Symmetry imposed | C2 | C2 | C2 | C2 | C1 |
| Overall resol. (Å) | | | | | |
| FSC=0.143 (masked) | 3.25 | 2.94 | 3.19 | 3.20 | 3.69 |
| Map sharpening B-factor (Å$^2$) | -168.4 | -141.6 | -152.4 | -167.5 | -122.1 |
| Local resol. rang.(Å) | 2.869-5.503 | 2.685-5.164 | 2.889-6.583 | 2.896-47.60 | 3.275-10.43 |
| **Model Refinement** | | | | | |
| Software | Phenix (v1.20.1-4487) | Phenix (v1.20.1-4487) | Phenix (v1.20.1-4487) | Phenix (v1.20.1-4487) | Phenix (v1.20.1-4487) |
| Non-hydrogen atoms | 5,054 | 5,050 | 5,032 | 5,028 | 10,174 |
| Protein residues | 662 | 662 | 658 | 658 | 1,333 |
| Ligands | 2 | 2 | 2 | 2 | 4 |
| *Av. B factors (Å$^2$)* | | | | | |
| Protein | 184.06 | 171.86 | 183.73 | 200.13 | 190.77 |
| Ligands | 253.47 | 215.50 | 216.35 | 55.94 | 213.07 |
| *R.M.S. deviations* | | | | | |
| Bond length (Å) | 0.002 | 0.002 | 0.002 | 0.002 | 0.002 |
| Bond angle (°) | 0.429 | 0.405 | 0.455 | 0.491 | 0.487 |
| *Ramachandran statistics (%)* | | | | | |
| Outliers | 0.00 | 0.00 | 0.00 | 0.00 | 0.00 |
| Allowed | 5.17 | 2.74 | 4.89 | 6.12 | 4.15 |
| Favored | 94.83 | 97.26 | 95.11 | 93.88 | 95.85 |
| MolProbity score | 1.56 | 1.27 | 1.61 | 1.83 | 1.83 |
| *Model vs. Map FSC* | | | | | |
| FSC=0.5 (masked,Å) | 3.5 | 3.3 | 3.6 | 4.2 | 4.1 |
